# Linking Codon- and Protein-Level Mutation Scores to Population Genetics Reveals Heterogeneous Selection Efficiency Across *Escherichia coli* Lineages

**DOI:** 10.64898/2026.03.16.711857

**Authors:** M. Mischler, L. Vigué, G. Croce, M. Weigt, O. Tenaillon

**Affiliations:** Université Paris Cité, Inserm, Institut Cochin, F-75014, Paris France; Department of Oncology UNIL CHUV, Ludwig Institute for Cancer Research, University of Lausanne, Lausanne, Switzerland; Sorbonne Université, CNRS, Dept. of Computational, Quantitative and Synthetic Biology UMR 7238, Paris, France

**Keywords:** Direct Coupling Analysis, population genetics, *Escherichia coli*, effective population size, Distribution of Fitness Effects, codon usage, Poisson Random Field

## Abstract

Quantifying the selective effects of individual mutations is essential to understand how their population-wise frequencies evolve under natural selection and genetic drift. Large genomic datasets provide a real-life experiment that we exploit to characterize the efficiency of selection across different mutations types and populations. Using Direct Coupling Analysis, a model from statistical physics, we derive protein-informed scores for individual non-synonymous mutations identified in 81,440 *Escherichia coli* genomes. We show that these scores act as a latent variable capturing the probability that a mutation is beneficial, neutral, or mildly to highly deleterious. We contribute to the debate on the importance of synonymous mutations by demonstrating that their selection intensities span a single order of magnitude in the *E. coli* species, whereas non-synonymous mutations span six orders of magnitude. We further relate selection efficiency to genetic drift, defined as the inverse of population size, and to ecological lifestyle, and we identify a 10,000-fold reduction in selection efficiency between the entire *E. coli* species and its most pathogenic populations. Together, these results highlight how population genetics and protein variant fitness predictors inform one another: variation in selection efficiency is associated with shifts in the distribution of mutation scores, and population genetics data provide a benchmark to assess the accuracy of these scores.

**Graphical abstract:** Schematic representation of the analysis of polymorphism in 81,440 Escherichia coli genomes. 458,443 polymorphic codon sites were identified and oriented using homologous sequences from closely related species. Mutations can be classified as synonymous or non-synonymous based on whether they alter the amino-acid sequence encoded, and real-valued scores predictive of fitness effects can be attributed to mutations within each of these classes. Codon scores reflect the global codon usage preference within the E. coli genome. DCA scores capture position- and amino-acid-specific preference as well as epistatic constraints and are obtained for each protein from a set of distantly related homologous sequences. Coupled with the abundance of polymorphic sites within different E. coli subpopulations, these different polymorphism classifications allow to precisely compare the intensity of selection between different types of mutations and across populations with distinct lifestyles, illustrated here by their pathogenic power.

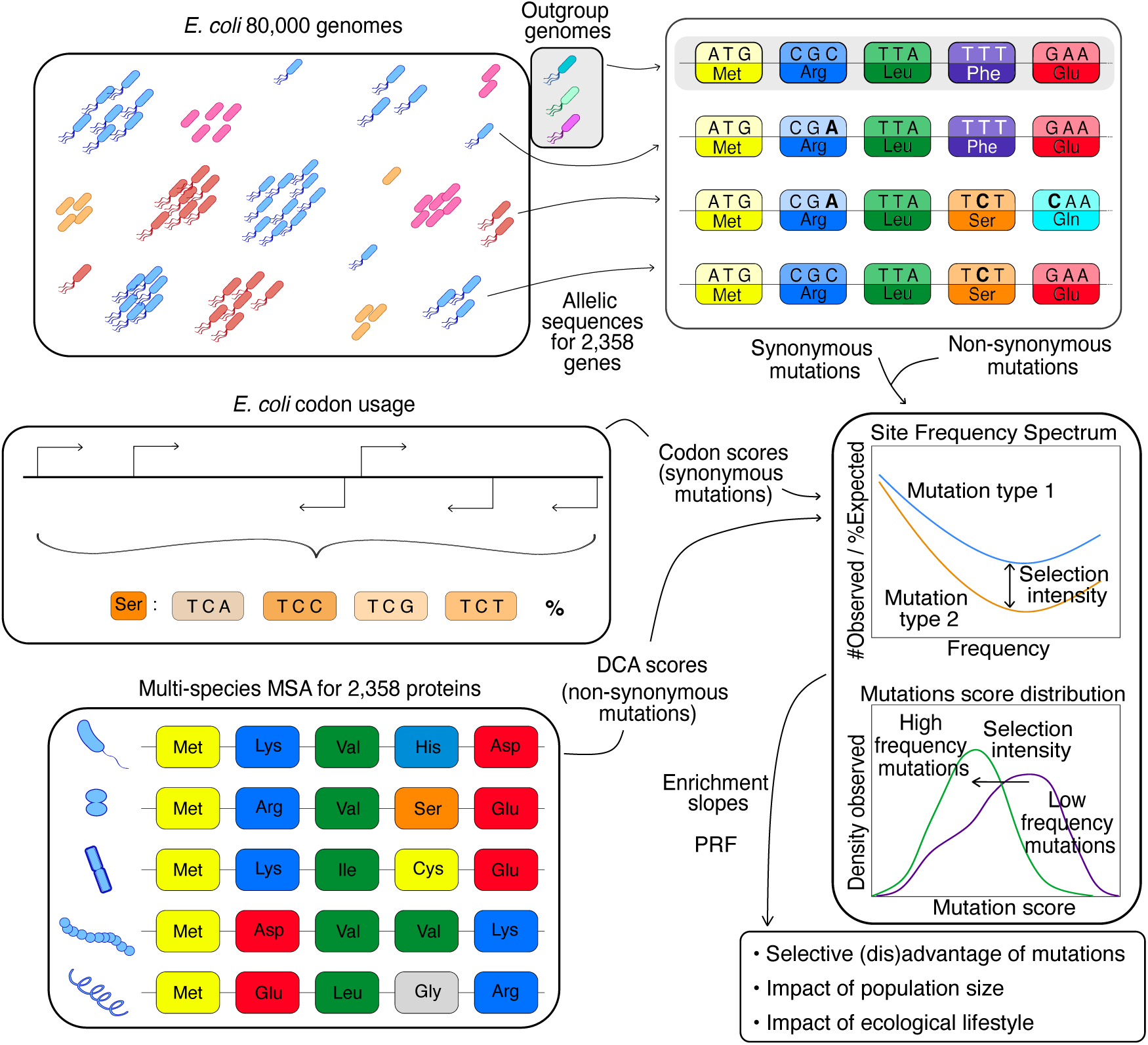

## Introduction

Genetic sequences in modern organisms act as records of evolutionary history, shaped by past selective pressures and demographic processes. Detecting these processes directly from genetic data has become feasible with the advent of high-throughput sequencing^1,2^. However, genomic sequencing has dramatically increased over the last few years, reaching hundreds of thousands of genomes for some species of medical interest. This wealth of genomes allows not only to quantitatively refine previous results on natural selection within bacterial species, but also to investigate the dynamic aspects of natural selection by comparing different subpopulations across distinct ecological niches, bridging the gap between long-term diversification and short-term population genetics.

Genetic variation within populations is often measured using polymorphic sites, which carry different alleles between otherwise similar genetic sequences. These sites can provide key information to population geneticists, such as the frequency of the mutated alleles in the sampled population and their functional impact. In protein-coding genes, which make up around 90% of bacterial genomes, mutations are either synonymous or non-synonymous. The former leave protein sequences unchanged and are typically assumed neutral, while the latter alter amino acids and are more likely to influence fitness. Most non-synonymous mutations are deleterious, but their effect vary widely. This variation depends on the amino-acid change, its position in the protein, and the protein’s importance to the organism. To describe these effects, population geneticists use a selection coefficient, *s*, which measures the fitness change of a mutation relative to the wildtype. Beneficial mutations have positive *s*, and deleterious mutations have negative *s*. Across the genome, mutation effects are commonly modeled as independent drawings from a Distribution of Fitness Effects (DFE), describing the variability in magnitude and sign of *s*.

A majority of mutations occurring by chance are expected to be non-synonymous^3^ and deleterious^4^. However, polymorphisms detected in small samples of natural populations are enriched for synonymous mutations^5^ as natural selection removes the most harmful variants, a process known as purifying selection. The same pattern is observed for the mutations fixed between two species^5^. This enrichment is quantified by the ratio of observed non-synonymous to synonymous substitutions, corrected for their respective mutation rates, known as *Ka/Ks* or *dN/dS*. Due to the finite size of populations, mutation frequencies also evolve partly in a stochastic manner, through genetic drift. Drift allows some mildly deleterious mutations to persist or even reach high frequencies. As a result, selection is not absolute and becomes more efficient in larger populations^6^. *Ka/Ks* is thus expected to be negatively correlated with a population’s effective size *N*_*e*_, which is often estimated from the population’s genetic diversity. This correlation has been confirmed by looking at the fixed mutations between pairs of related eukaryotic species across many taxa^7^. Here, we test whether the same relationship exists among bacterial subpopulations within a single species, in particular when these subpopulations rely on distinct ecological strategies for their propagation.

*Ka/Ks* can be used to contrast genes or lineages under different selective constraints in experimental or natural contexts. However, with sufficiently large samples, a more informative approach compares non-synonymous and synonymous polymorphisms across each allele frequencies^8–11^: the Site Frequency Spectrum (SFS) of putatively non-neutral mutations is compared to the SFS of putatively neutral mutations. As mutations with more deleterious effects are filtered at lower frequencies by purifying selection, comparing these two SFS provides information on the DFE. The absence of some non-synonymous mutations at the lowest observable frequencies corresponds to mutations that are so deleterious that they are eliminated before they can be sampled. The larger the sample, the smaller this missing fraction should be. Mutations with intermediate deleterious effects should start being depleted at corresponding intermediate frequencies *q*∼1/*N*_*e*_|*s*|^12^. Non-synonymous mutations that reach high frequencies are expected to be effectively neutral or beneficial. This framework allows to infer the global shape of the DFE from the collective behavior of non-synonymous mutations. However, all these mutations are treated equivalently, regardless of the protein in which they occur or the specific amino-acid change they cause. Yet these properties could provide valuable information about which region of the DFE a given mutation is likely to originate from.

Instead of relying on a single binary classification, a different approach consists in modeling the impact of non-synonymous mutations at the protein level. Although some population genetics studies have used the chemical properties of amino acids to classify substitutions into ‘radical’ and ‘conservative’ classes in a spirit similar to *Ka/Ks*^13^, more advanced and data-driven computational tools were initially developed in the field of protein structure prediction before being applied more recently in evolutionary and population genetics.

The proteins observed in nature today result from an evolutionary process that diversified their sequences while preserving their function through natural selection. This has led to families of homologous proteins – sequences that vary widely across species but share highly similar biological functions and folded structures. Building on the accumulation of sequence data in public databases, families of diverged homologs were identified for thousands of proteins, frequently represented via multiple-sequence alignments (MSAs). These MSAs encapsulate a historical and functional narrative of protein evolution, which computational tools can decode. The development of direct coupling analysis (DCA)^14^ has enabled the estimation of a “statistical energy” (or negative log probability) of a sequence as a function of two components: a position-specific conservation score, reflecting how suitable a given amino acid is at a particular position in this protein family, and a sum of epistatic coupling scores, indicating how well that amino acid interacts with others at different sites. Originally developed to identify residue interactions in MSAs^14^, this statistical-learning approach has also proven useful for the generation of artificial functional homologs^15^ and for the study of mutational effects, both at the single protein level^16–18^ and at the genome level^19,20^, in experimental evolution^19^ and natural populations^17,20^ alike. Thus, DCA provides a protein-customized prediction for the selective effect of any non-synonymous mutation. Keeping in mind that the same DCA scores values may reflect different selection coefficients owing to differences in functional importance and selective pressure among proteins, we can confront DCA-based predictions to a polymorphism dataset, ideally spanning species-wide diversity.

The bacterium *Escherichia coli* serves as an exemplary system for this purpose due to its extensive sequencing coverage and well-characterized population structure. Indeed, *E. coli* is a commensal of the gut of vertebrates used as a fecal contaminant marker in environmental science^21^, but also a versatile pathogen involved in both extra- and intra-intestinal disease such as diarrhea^22^, hemorrhagic colitis^22^, hemolytic uremic syndrome^22^, dysentery^23^, urinary track infection^22^, bloodstream infection^22^ and neonatal meningitidis^22^. It has gained increasing medical relevance in the recent decades due to the emergence of antibiotics-resistant clones that limit the treatment arsenal and has made the species the top killing bacterial species associated with resistance^24^. The diversity of lifestyle in *E. coli* results from its genomic diversity, which can be organized in diverse clonal clusters using several typing methods such as serotyping or Sequence Typing (ST), or more advanced genome-wide methods such as hierarchical clustering^25^ and genetic distance clustering – which we use in this article. Genomic methods have definitively anchored the intracellular pathogens known as *Shigella* within the *E. coli* species, from which they repeatedly evolved^26^. However, due to its medical importance, the genus name *Shigella* and its ‘species’ *S. dysenteriae, S. boydii, S. flexneri* and *S. sonnei* are still in use, although they were defined in the early 20^th^ century based on biochemical criteria that only partially overlap with phylogenetic relations. We use more recent denomination based on genetic evidence to refer to the different *Shigella* clusters^26,27^. *Shigella* representative genomes from different clusters have already been described as coming from populations of reduced sizes, based on single genome metrics such as a high number of Insertion Sequence (IS) and a high proportion of non-synonymous to synonymous substitutions during their evolution from the rest of *E. coli*^2,28^. However comprehensive population-level comparison between *Shigella* clusters and other pathogenic and non-pathogenic *E. coli* clusters is still lacking to our knowledge.

Beyond its clinical and environmental importance, *E. coli* has been a cornerstone of molecular biology, with much of its physiology thoroughly studied. This extensive understanding has made it an ideal model for evolutionary research, exemplified by pioneering studies linking DNA resequencing to adaptation^29,30^, including study of synonymous/non-synonymous fixed mutations^31^ as well as the first use of DCA scores to estimate selection at the genome level^19^. This combination of evolutionary, medical, and experimental relevance makes *E. coli* a particularly suitable model for studying the impact of natural selection on genetic diversity. In this work, we leverage over 80,000 *E. coli* genomes and integrate codon- and protein-level mutation scores with population-genetic analyses to quantify selection pressure across *E. coli* lineages (Graphical Abstract).

## Results

### 1. Introducing genetic polymorphism in the *E. coli* database

We aim at finding the largest possible number of polymorphisms present in the complete *E. coli* species and to orient them by determining, for each site, which codon represents the most probable ancestral state. Our genomic dataset is a complete download of Enterobase^32,33^ in February 2019 and is constituted of 81,440 *E. coli* and *Shigella* genomes. We identified close to 350,000 different genes (Fig. 1A, see Methods) and decided to focus on the core genome, defined as the set of genes present in almost the entire species (>95% of the genomes) and almost never duplicated (<1% of the genomes). As *E. coli* genomes are highly dynamic with quick gain and loss of genes^2^, the fact that these genes are core suggest that they are necessary for *E. coli* in at least some environment it regularly experiences, and that they are under strong purifying selection. We focused on the core genes that were 1) less than 800 amino-acid long, 2) for which we could find at least one homologous sequence among three closely related species (‘outgroups’) and 3) a MSA comprising >200 distant homologs could be gathered. We obtained a set of 2,358 genes, covering altogether >2,000,000 bases and around 40% of a typical *E. coli* genome. We then aligned all alleles found in the database and gathered codon sites that displayed polymorphism within the 81,440 genomes (90.6% of all codon sites). We retained the sites for which 1) the two most abundant codons were present in more than 95% of the genomes (89.1% of all sites) and 2) one and only one of the observed alleles was present in the set of outgroup codons (66.3% of all sites). We were thus able to orient 458,443 mutations (Supplementary Fig. 1).

**Fig. 1:**
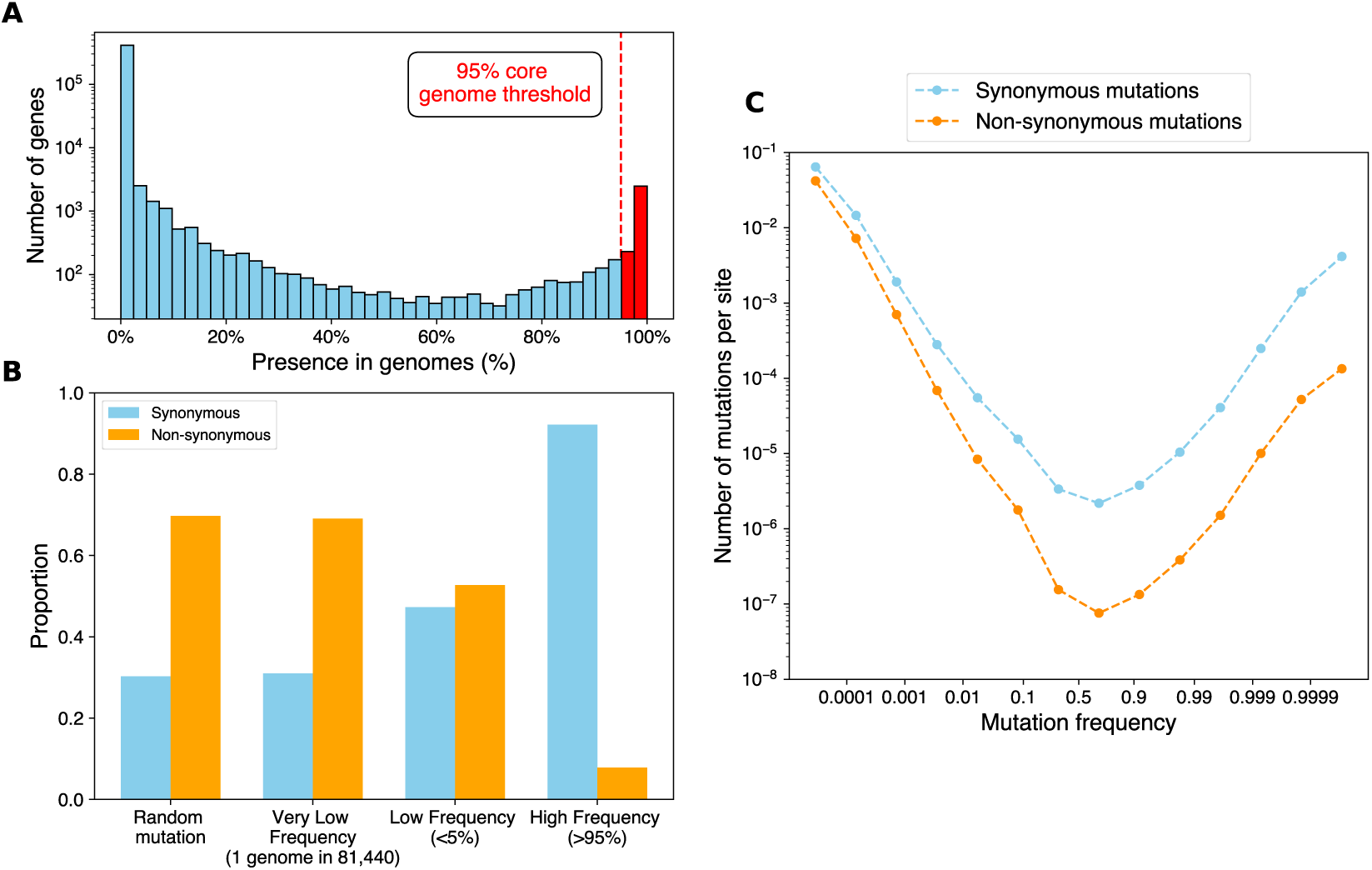
Core genome definition and segregation of synonymous and non-synonymous mutations in the 81,440 genomes database. **A.** Histogram of the presence of genes among the 81,440 genomes. A threshold of 95% presence was used to select genes under strong purifying selection. **B.** Proportion of synonymous and non-synonymous mutations at different frequencies in the database within the 2,358 selected core genes. The proportion for random mutations (30.3%-69.7%) was computed using the base-to-base mutation biases inferred from low frequency mutations. **C.** Site frequency spectrum of synonymous and non-synonymous mutation, represented on a logit scale with binned frequency. The number of mutations per site corresponds to the number of mutations found within a certain frequency bin, divided by the size of the bin, the length of the total alignment (in codons), the proportion of successfully oriented polymorphisms among all polymorphic sites and the probability of a new mutation being from this category (synonymous or non-synonymous).

Having identified the mutations frequencies’ as well as their ancestral and mutated state, we could now use different proxies to estimate their effect. A small half (45.9%) of these mutations are non-synonymous and the rest is synonymous. However, this proportion is highly dependent on the frequency at which the mutations are found (Fig. 1B): for extremely low frequency (mutations carried by only 1 genome among the 81,440, that is ∼0.0012% frequency), the proportion is very close to the chance of obtaining a synonymous mutation when mutating a base randomly, which is around 30%. However, this proportion increases with frequency: mutations present in up to 5% of the genomes are in equal proportions synonymous or non-synonymous, and mutations close to fixation (>95% frequency) are overwhelmingly synonymous (89.3%). The progressive enrichment of synonymous mutations over non-synonymous mutations can be visualized by comparing the two mutation types’ SFS on a log-logit-scaled axis (Fig. 1C). The SFS are scaled by the total alignment length and the probability of a new random mutations falling in the mutation class (synonymous vs non-synonymous). Due to this scaling, the two SFS start almost at the same value (0.04 and 0.064), which is very close to the species’ diversity (*θ*_W_ = *S*⁄log (*n*_*sample*_) = 0.08). The decrease in the number of synonymous mutations with frequency is an illustration of genetic drift: for the low frequencies *q* ≤ 0.1, the synonymous SFS decreases with the inverse of *q*, as expected under simple neutral population genetics models. The swifter decrease observed for non-synonymous mutations is due to the combined effect of drift and purifying selection. At higher frequencies (*q* ≥ 0.5), both SFS show an increase not expected under the most simple neutral population genetics model but widespread in SFS from natural population^34^, which could reflect the impact of changes in population size, population structures, finite recombination or could be a consequence of the sampling procedure, among others^34^.

### 2. DCA distributions and enrichment slopes

With this evidence for the presence of purifying selection in mind, we would like to investigate if the non-synonymous mutations that do reach high frequencies have a DCA signature distinguishing them from other non-synonymous mutations.

As DCA is a continuous score, the joint distribution of non-synonymous mutations DCA score and frequency in the population (Fig. 2A) can be explored in two manners: by looking how the distribution of scores shifts with increasing frequencies (front view) or by regrouping mutations with similar DCA scores together and comparing the SFS of each of these classes (side view).

**Fig. 2:**
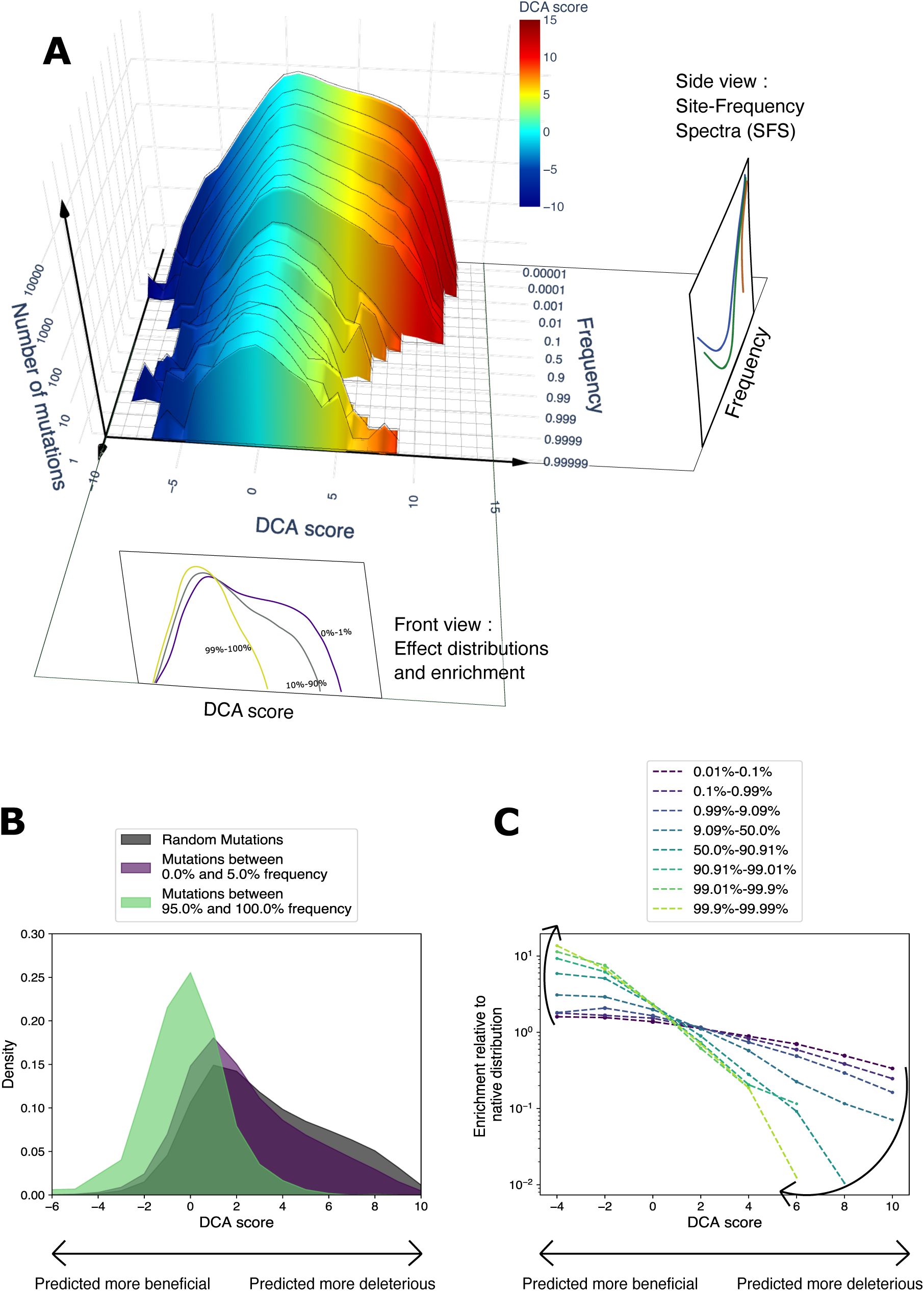
Distribution of the DCA scores and frequency within the database of oriented non-synonymous mutations within the entire database. **A.** Joint distribution of DCA scores and frequencies. The z-axis represents the absolute number of mutations in the corresponding DCA bin and frequency bin. The SFS can be seen as the marginals for different values of DCA scores conditioned by the mutations being in this range of DCA score. The distributions of effects correspond to the marginals for mutations observed at increasing frequencies. **B.** Distribution of DCA scores at different frequencies. Black: distribution of scores of new non-synonymous mutations, occurring at random and prior to any selection. Purple: observed distribution of scores for mutations found below 5% frequency. Green: observed distribution of scores for mutations found above 95% frequency. **C.** Ratio of the observed distribution of scores by the distribution expected for random mutations, for increasing frequencies of observed mutations. The ratio appears to be exponentially decreasing with DCA scores, and more strongly at higher frequencies.

DCA scores represent a difference in statistical energy, so mutations with negative scores are predicted beneficial – they have a higher probability under the DCA model – while mutations with a positive score are predicted deleterious. As DCA models epistasis, the score of a single amino-acid substitution depends on the rest of the protein sequence (its background). Although the score depends strongly on the background, it does so globally, so the mutational effect of a given substitution shifts little between close sequences (intra-species) and this shift only becomes significant for diverged sequences, typically from distant species^20^. Thus, we computed all DCA scores in the context of the *E. coli* protein consensus. The score distribution of all randomly occurring mutations is strongly biased for positive values (Fig. 2B), which is consistent with the expectation that most randomly occurring non-synonymous mutations are deleterious. However, mutations effectively observed at low frequency (0% to 5%) have a score distribution still biased for positive values, but much less than random mutations: it is only slightly shifted towards negative DCA scores, that correspond to less deleterious mutations. Mutations that are close to reaching fixation (95% to 100% frequency) have a score distribution symmetrical and centered around 0. An interesting metric to quantify the intensity of this shift is the ratio of score distribution by the distribution of random scores (Fig. 2C). On a log scale, this ratio is linearly decreasing with DCA with a slope whose steepness increases when considering mutations segregating at higher frequencies.

An important prediction from population genetics theory is that selection is more efficient in larger populations. We can test this prediction by comparing the shift in DCA distributions in different *E. coli* sub-populations. Indeed, the 81,440 genomes can be divided into 579 distinct clusters based on their genetic distance (see Methods). We chose a neighborhood threshold of 0.5% different bases in the core genome alignment to define clusters, as the distribution of pairwise distances in the database shows a gap around this value (Supplementary Fig. 2) and this method is currently used to identify cryptic subspecies^35^. As a comparison, the average genetic distance between human and chimpanzee is slightly higher than 1%^36^.

The clusters found with this method align with previous classifications, in particular for major epidemiologic lineages (Fig. 3A and Supplementary Fig. 3). Most *E. coli* lineages can become opportunistic pathogens and cause extra-intestinal infection, with variation between lineages viewed rather as a coincidental by-product of adaptation to *E. coli*’s main niche, the mammalian gut, than as a result of selection for virulence^37^. Unlike these commensal / extra-intestinal pathogenic *E. coli* (ExPEC), some other lineages will systematically cause intra-intestinal infection. These intra-intestinal pathogenic *E. coli* (InPEC) can be divided into 3 classes: STEC (Shiga toxin-producing *E. coli*) including EHEC (enterohemorrhagic *E. coli*) that closely adhere to the surface of human intestinal cells (enterocytes) and produce toxins, *Shigella* lineages that can invade human cells and have a reduced genome, and finally entero-invasive *E. coli* (EIEC) that are similar to *Shigella* but with a less reduced genome (Fig. 3B). Each of these pathotypes has evolved several times from non-pathogenic *E. coli,* mainly driven by the horizontal transfer of virulence genes as well as the repeated loss of some genes in the case of *Shigella* and EIEC^38^. In order to correct for different sample sizes for different clusters, we limited our analysis to clusters with more than 110 genomes and subsampled each one 10 times down to 100 genomes. Clusters were classified into the 4 pathotypes (see Methods). When comparing with the total *E. coli* population, a set of representatives from each cluster was selected (‘Diversity sample’, 718 genomes). For each of these populations, we computed the slope of the shift in DCA distribution observed at a frequency higher than 50% (similar to Fig. 2C).

**Fig. 3:**
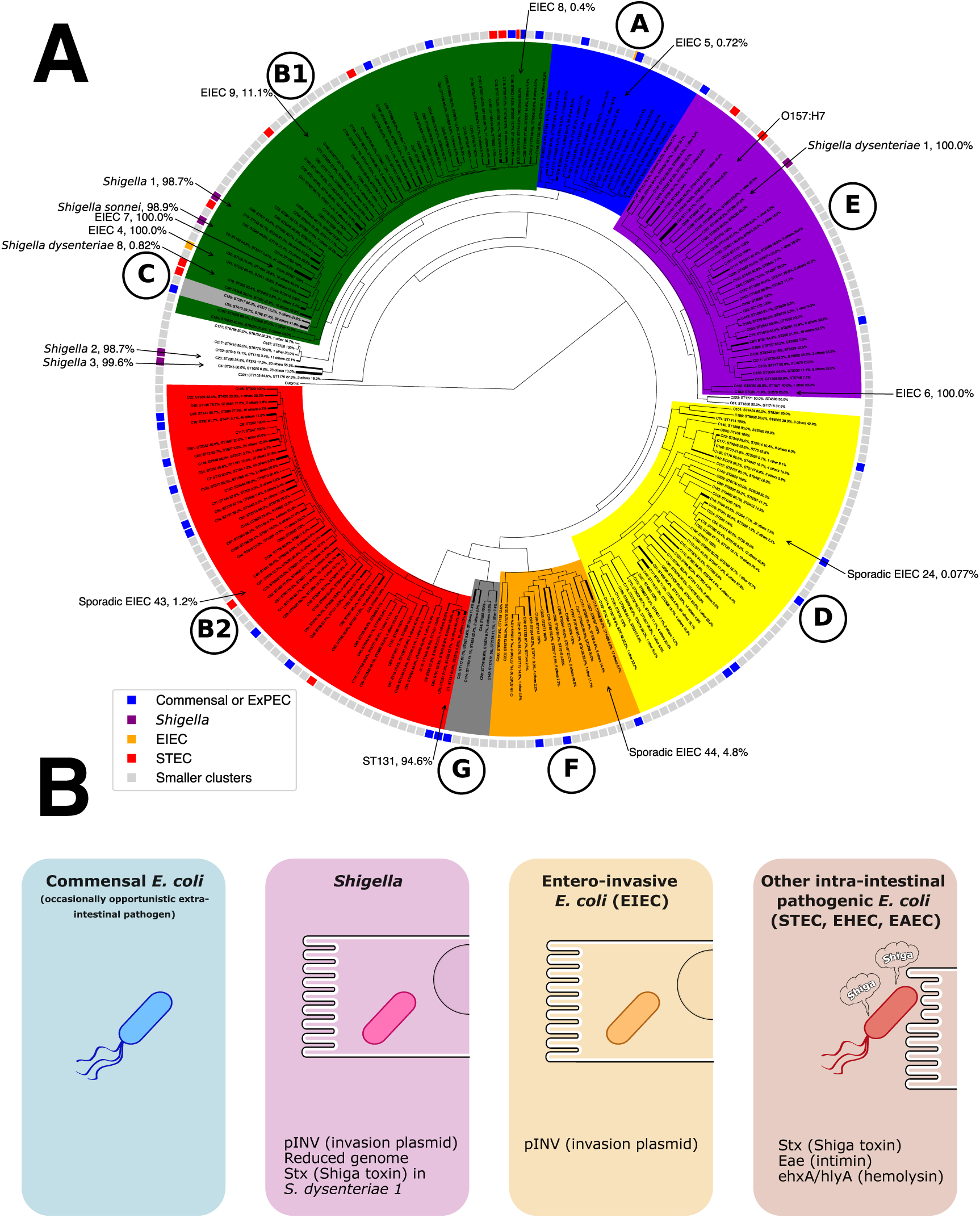
Phylogeny of the main clusters and schematic description of the 4 lifestyles studied here. **A.** Phylogenetic tree of the ancestors of each one of the 240 main clusters (see Methods), rooted with *E. fergusonii* ATCC 35469T. Phylogroups A to G are represented as colored sections and the outer rim indicates the lifestyle of the majority of the cluster for clusters with more than 110 isolates. Note that the two largest clusters, cluster 3 (B1) and cluster 8 (A), encompass several lifestyles with a small minority of EIEC in both and a significant minority of STEC in cluster 3 (see Supplementary Table S2) **B.** Representation of 4 *E. coli* lifestyles: the commensal and/or environmental strains, that can occasionally cause extra-intestinal infections (ExPEC, extra-intestinal pathogenic *E. coli*); the specialized intracellular pathogen *Shigella* that can infect enterocytes due to its invasion plasmid pINV; the entero-invasive *E. coli* (EIEC) that acquired pINV more recently than *Shigella* and can still thrive outside human cells; and the Shiga-toxin producing *E. coli* (STEC), which include enterohemorrhagic *E. coli* (EHEC), that attach to the surface of enterocytes and produce toxins against their host.

*Shigella* clusters show the less abrupt shift in their DCA distribution (Fig. 4A), while the *E. coli* cluster representatives show the strongest one (Fig. 4D). This confirms that the globally reduced selection in *Shigella* found by earlier genomic studies^28,39^ also translates into a lower selection intensity at the population level and into a lower ability to get rid of the most deleterious non-synonymous mutations. All others pathotypes are significantly more depleted in highly positive DCA scores than *Shigella*, with commensals showing the strongest shift after entire *E. coli* species (Fig. 4B-D), suggesting that the population reduction observed for *Shigella* is also present in other InPEC clusters, although less strong (Fig. 4F).

**Fig. 4:**
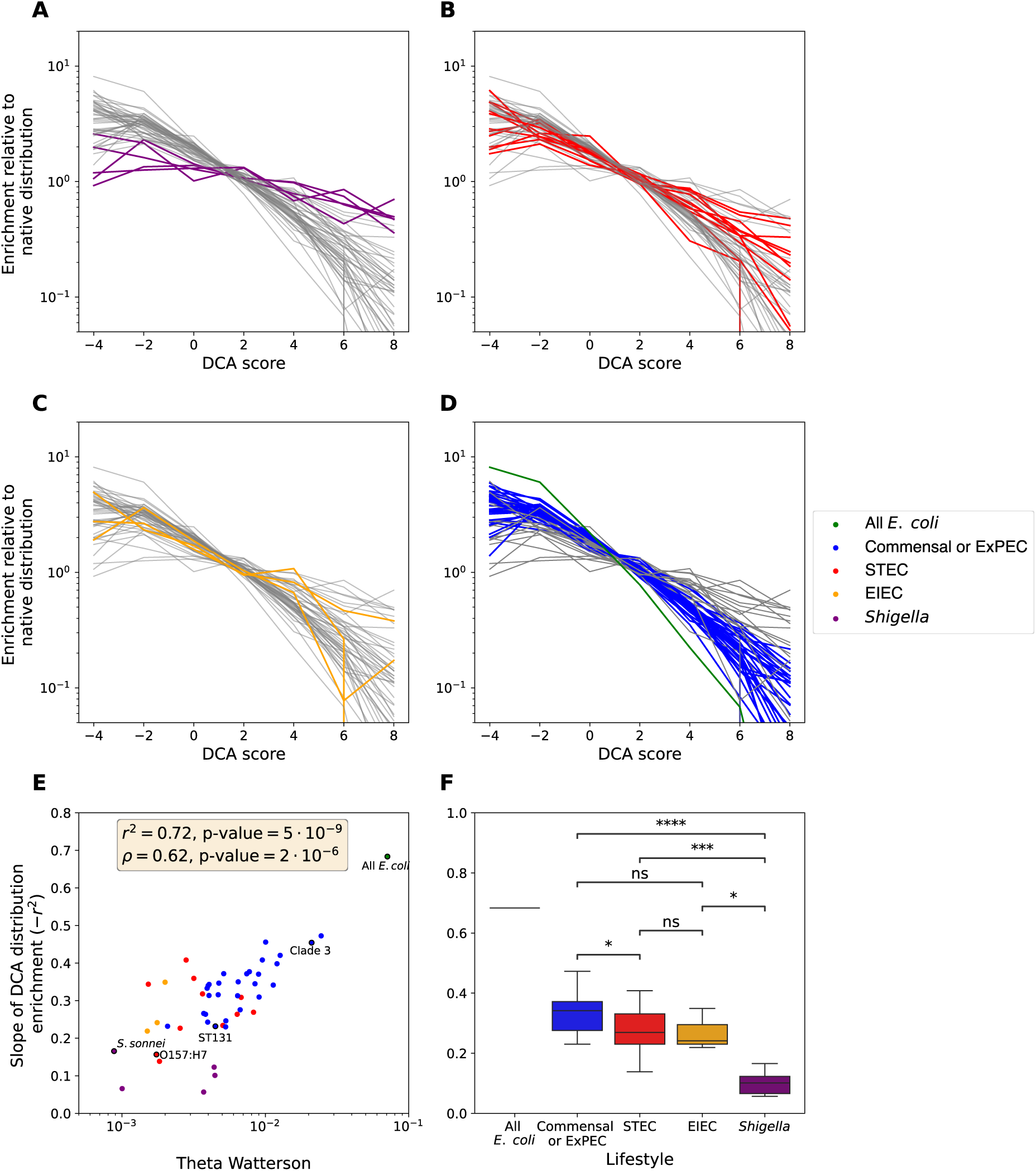
DCA enrichment slopes for the 49 main clusters. **A. – D.** Enrichment curves with the 4 pathotypes successively highlighted. **E.** Linear correlation between the intra-cluster diversity and its enrichment slope. **F.** Enrichment slope grouped by cluster’s lifestyle. Welch’s t-test independent samples was realized between each pair of lifestyles. *: p-value ≤ 0.05; **: p-value ≤ 0.01; ***: p-value ≤ 10^−3^; ****: p-value ≤ 10^−4^. Remind that we only have 3 EIEC populations.

Clusters’ enrichment slope correlates with their genetic diversity measured by the theta Watterson estimator *θ*_W_(Pearson *r*^2^ = 0.68 with p-value of 8 ⋅ 10^−8^, Spearman *ρ* = 0.62 with p-value of 2 ⋅ 10^−6^). Population geneticists often model species diversity by an effective population size, *N*_*e*_, that corresponds to the size an ideal Wright-Fisher population would need to have to display the genetic diversity observed. Although *N*_*e*_ is usually estimated directly from genetic diversity, it is also related to the efficiency with which deleterious mutations are purged from a population. Population geneticist often compare *Ka/Ks* or related metrics to genetic diversity to validate this population size dependency, and we show that this effect can also be captured by the slope of DCA scores distribution shift (Fig. 4E). Thus, DCA enrichment slopes provide a semi-quantitative proxy for individual mutations with a statistical behavior coherent with what is expected from the selective coefficient of a mutation. However, the exact relation between DCA scores and the distribution of fitness might be more complex and we need to introduce a population genetics model to investigate this relation. Before applying this model on the DCA scores of non-synonymous mutations, we first validate it on a less complex type of mutations.

### 3. Introducing a selection model for Codon usage

When genetic polymorphism sequencing data became available in the 1990s, they were reduced to a few genes sequenced in a few individuals, motivating the development of a probabilistic framework called the Poisson Random Field approach (PRF)^40^ able to deal with the uncertainty arising from these relatively small samples. It also adapted the continuous diffusion equations used in population genetics models to discrete genetic data by explicitly modeling the sampling process (see Methods). Since its first applications to sequencing data^8^, population geneticists have contrasted the discrete SFS of non-synonymous mutations to the one of synonymous mutations taken as a neutral reference, an assumption often left unchecked. As recent experimental works have reported synonymous mutations with selective effects in microbial populations^41,42^, we first investigate if the well-known impact of synonymous mutations deriving from preferential codon usage can leave traces in *E. coli* polymorphisms.

Preference for each codon can be estimated from their global usage in the consensus coding sequences of core genes: for instance, the 6-fold degenerated amino acid Leucine is encoded by CTG in 53.6% of the core sites and by CTA in only 2.9%. Each synonymous mutation can be attributed a score defined as the logarithm of the ratio of the usage of the ancestral to the mutated codon:

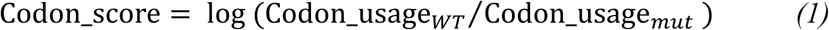

As for DCA scores, negative scores are characteristic of a mutation from a less used codon to a more used one and should be beneficial, while positive scores reflect a change toward a rarer codon and should be deleterious. The distribution of codon scores can be compared for mutations found at different frequencies (Fig. 5A-B). As the random distribution is directly related to the mutation rate of individual bases and codons, its estimates can be influenced by lineage-specific mutation biases or other selective constraints, in particular for extreme codon score values which depends on a few codon substitutions. Thus, instead of relying on base-to-base mutation rates or single-genome mutations which could be influenced by sequencing biases, we use the mutations present in between 3 and 5 genomes among 81,440 to estimate the distribution of codon scores arising by random mutations. When looking at mutations found at low frequencies, we obtain a flat distribution ratio and overlapping SFS (Fig. 5B-C). At these low frequencies, the favorable or unfavorable synonymous mutations have not yet reached the threshold to become visible by natural selection and are all segregating neutrally: due to their low effects, the transition between these two behaviors happens at relatively high frequency and can be explicitly mapped.

**Fig. 5:**
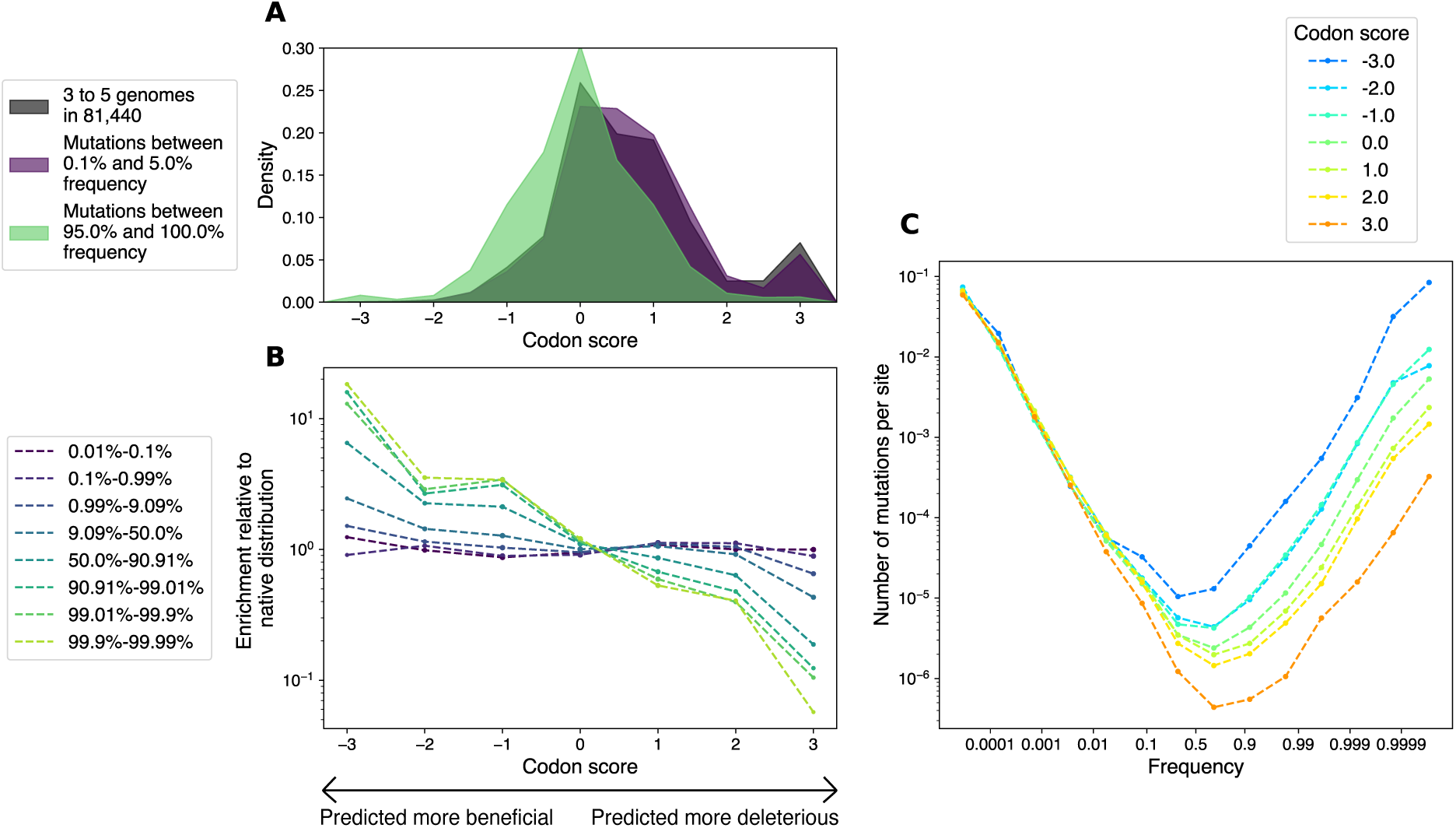
Distributions of Codon scores and SFS of synonymous mutations. **A.** Progressive shift in the distribution of Codon scores of synonymous mutations observed at increasing frequency. **B.** Enrichment slopes of Codon scores become steeper as frequency increases. **C.** SFS of synonymous mutations with different Codon scores. Each SFS is scaled by the probability that a new mutation appears with a Codon score in the corresponding range of values.

Next, we investigate how codon usage-related selection can be quantified. To do so, we assume that synonymous mutations with a codon score close to 0 are neutral as they don’t result in an important change in codon usage. Selection intensities on other codon score values can then be estimated by comparison to this neutral reference. This is similar in spirit to the PRF approach which uses synonymous mutations as a reference to detect selection on non-synonymous mutations, except that we are looking at intra-synonymous selection here. However, the PRF approach is adapted to small or medium sample sizes and treats mutation frequency as a discrete variable. With a sample of 81,440 genomes, it is not unreasonable to treat frequency as a continuous variable and directly infer the selection intensity by fitting the observed SFS against the theoretical one. With this ‘curve-fitting’ method the goodness of fit can be assessed visually by comparing the observed SFS to the best-fitting inferred curve (Fig. 6A). To validate this new method, we compare it to the PRF method used on the diversity-enriched subsample of 718 *E. coli –* a sample size small enough for this approach to be effective.

**Fig. 6:**
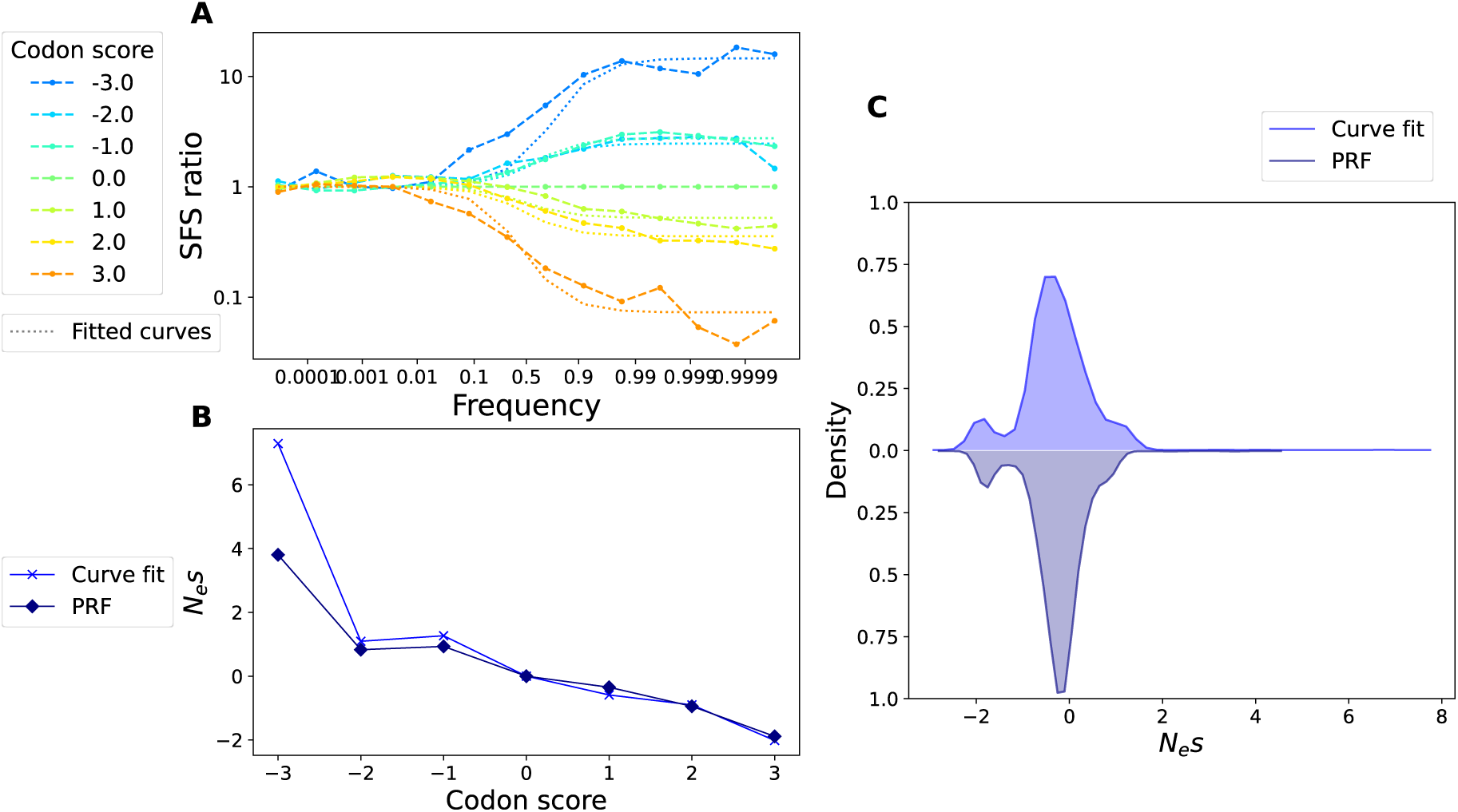
DFE inference for synonymous mutations. **A.** Observed SFS ratio and best-fitting SFS ratio (dotted lines) with the curve-fit single effect method for synonymous mutations with different Codon scores. The SFS ratio is computed over the entire *E. coli* dataset. **B.** Selective intensity *N*_*e*_*s* inferred for each DCA score value with the curve-fit method (on 81,440 genomes) and the PRF method (on the representative set of 718 genomes). **C.** Distribution of selection intensities obtained with the two methods.

Both methods estimate the ratio of selection over drift, called selection intensity:

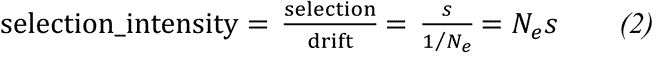

The selection intensity is inferred independently for each range of Codon score based on how mutations with scores in this range are collectively enriched over neutral mutations. Both methods lead to weak selective intensities for codon usage, with |*N*_*e*_*s*| always smaller than 10 (Fig. 6B) and often smaller than 1 (Fig. 6C), for positively selected or negatively selected codon usage scores alike. Although the associated selection efficiencies are rather small, selection for codon usage is detectable from polymorphism alone within the *E. coli* species. Remarkably, we could fit each SFS ratio with a single effect for each Codon score values and obtain a good fit, while approaches that don’t rely on such precise scores have to make the assumption of a distribution of effects *while* fitting. In our case, we could determine the effect attributable to *each* mutation, and the distribution of effects we obtain is deterministically linked to the distribution of Codon scores. Unless explicitly mentioned, we use the synonymous mutations with a codon score close to 0 as a neutral reference in the rest of the analysis.

### 4. Extending to DCA scores with distribution of effects

We can now focus on DCA scores of non-synonymous mutations. We first compare the non-synonymous SFS of various DCA scores to that of the synonymous neutral reference (Fig. 7A and B) aforementioned. A first striking sight is the scale of the depletion of the mutations with the worst DCA scores, which reaches 100- and even 1000-fold compared to the less than 10-fold enrichment or depletion observed for the synonymous mutations. Moreover, the enrichment pattern is more complex and does not follow what is expected from a group of mutation with a unique selective intensity (Fig. 7B and Supplementary Fig. 4). In particular, most SFS are already depleted at the lowest observable mutation frequencies, which is characteristic of very strongly deleterious or lethal mutations. However, these SFS still reveal the presence of mutations up to intermediate frequency (10 to 50%), which is not compatible with a single strong effect. This pattern is coherent with a distinct distribution of fitness effects for each DCA score class, unlike what we obtained for Codon scores.

**Fig. 7:**
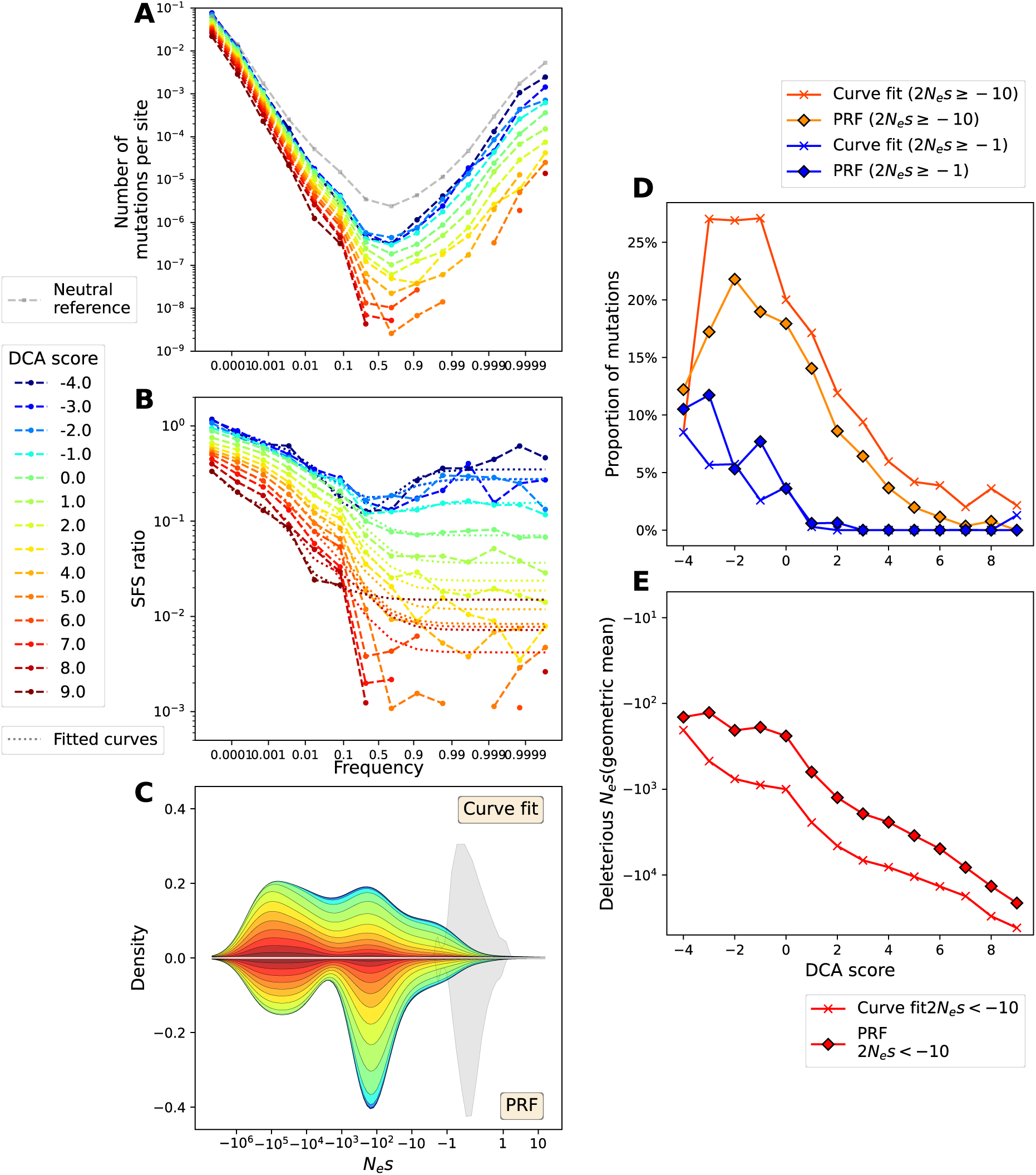
DFE inference for non-synonymous mutations for the entire *E. coli* species. **A.** SFS of non-synonymous mutations with different DCA scores. Each SFS is scaled by the probability that a new mutation appears with a DCA score in the corresponding range of values. The SFS of synonymous mutations with a Codon score between −0.5 and 0.5 (neutral reference) is depicted with grey dashes. **B.** Observed SFS ratio and best-fitting SFS ratio (dotted lines) with the curve-fit binned distribution method (see Methods). **C.** Distribution of selection intensities obtained with the curve-fit and the PRF methods, colored by DCA scores. Notice that the total area colored for a DCA score corresponds to its probability of appearance. The distribution of synonymous selection intensities is represented in grey, and scaled with the ratio of synonymous to non-synonymous probabilities of appearance. **D.** Proportion of mutations with a selective effect higher than −5 (weakly deleterious to beneficial, orange) or higher than −0.5 (effectively neutral to beneficial, blue), as a function of DCA scores. The quantity on the y-axis corresponds to the fraction of the mutations with these effects within all mutations with a DCA score in the corresponding range. **E.** Geometric mean of the remaining effects (selection intensity lower than −5) as a function of DCA score.

This is not unexpected, as non-synonymous mutations result in much more complex effects than synonymous mutations. DCA aims at predicting these effects by taking into consideration site-specific amino-acid preference and interactions with other amino-acids within the protein. However, proteins form complexes with other proteins and biomolecules. DCA models trained on single-protein MSAs do not capture these interactions. Even within the protein, DCA predictions may not be perfect due to the high number of parameters it needs to estimate (proportional to the square of the protein length). Finally, some core proteins are essential for immediate cell survival, whereas others can tolerate mildly deleterious mutations over short timescales. These mutations may not strongly affect short-term survival but can reduce long-term lineage fitness. DCA models are trained on highly divergent sequence data, so they reflect long-term evolutionary constraints. As a result, they may not fully capture short-term selective pressures that differ among proteins and shape the polymorphism patterns we observe^17^.

This uncertainty in the estimation of DCA score can however be taken into account in our fitting strategy by inferring a distribution of effects for each score value. As we have no a priori on the particular shape that distribution should have, we fit discretized DFEs with several bins covering each an order of magnitude of selection intensity, an approach widely used in recent population genetics studies^10,11,43^. We can pull these DFEs together to represent the total distribution of effects of non-synonymous mutations (Fig. 7C) obtained with the PRF and the curve-fitting methods. Both methods find a distribution spread across 6 orders of magnitudes, that dwarves the distribution of synonymous mutations in term of effect size. Both suggest an important proportion of extremely deleterious mutations, with *N*_*e*_*s* between 10^4^ and 10^6^, consisting mostly in mutations with positive DCA scores. These mutations can be considered lethal as they correspond to the mutations with high DCA scores unobserved at very low frequency (Fig. 7B and Fig. 2B).

DCA scores with negative effects display a non-monotonously decreasing SFS ratio, with an increase at higher frequencies. This is characteristic of beneficial mutations (Supplementary Fig. 4), so we also include a bin of mildly beneficial mutations for these DCA score values. In order to better visualize how DCA scores lead to different distributions of effects, we focus on the proportion of mutations with mild effects (−10 ≤ 2*N*_*e*_*s* ≤ 10) as well as the proportion of neutral to beneficial effects (−1 ≤ 2*N*_*e*_*s* ≤ 10) (Fig. 7D). Both proportions decrease with increasing DCA score, independently of the method used. In particular neutral and beneficial mutations only occur for negative or null DCA scores, and these scores also exhibit the highest proportion of weakly deleterious mutations, although DCA scores with small positive values (up to 3) also display a significant fraction (>5%) of such weakly deleterious effects. Additionally, we estimate the mean effect of intermediate and highly deleterious mutations (2 *N*_*e*_*s* < −10) (Fig. 7E), using the geometric mean so as not to overestimate the contribution of a few very deleterious mutations. It decreases exponentially with DCA score with both methods.

Having characterized the distribution of effects at the species-wide level, we can now record its global shape and infer how reduced its scale is in different sub-populations with varying genetic diversity and varying equivalent population sizes *N*_*e*_. Although *N*_*e*_ is often studied at the species level, the genetic clusters we identified are well-separated by a genetic distance of 0.5%. They tend to share more accessory genes within each cluster than across, and are thus likely more ecologically similar to *E. coli* from their own clusters than from different ones, resulting in an intra-cluster competition that naturally selects favorable mutations and deplete the most unfavorable ones.

### 5. Comparing selection efficiency across different clusters

In order not to bias our population sizes estimations, we subsampled 10 times all clusters down to 100 genomes and used the PRF method, better suited for this kind of sample size – as we did when working with DCA distribution enrichment slopes. We used all synonymous mutations as a neutral reference, as some clusters have a reduced genetic diversity. We first inferred the DFE shape using the samples representative of the total *E. coli* species (‘DivSampleEcoli’). We then estimated the scaling factor by which to multiply this species-wide DFE to best fit the SFS of the smaller, less diverse clusters. Indeed, the PRF method infer the distribution of the compound parameter *N*_*e*_*s*. As *N*_*e*_ is characteristic of the population, the intra-population variation is due to an underlying distribution of *s*, that we can consider constant for all *E. coli*. Thus, as a first approximation, the DFE in each cluster can be modeled as the species-wide distribution scaled by the ratio 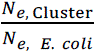.

This ratio measures the efficiency of purifying selection within each cluster relative to the species-wide level, which occur at longer timescales. Selection efficiency is lowest in *Shigella* clusters, where it represents only a fraction of the species-wide value (10^−4^ to 4 ⋅ 10^−3^ depending on the cluster). EIEC clusters and a majority of the STEC lineages are below 10% efficiency, while commensal/ExPEC clusters are between 10% and 80%. This result is coherent with what was previously found using DCA distribution ratio (Fig. 4) but with more power to discriminate between small population sizes, although it seems to saturate at higher population sizes with the most efficient cluster having a selection efficiency slightly below the one found for the whole species.

Having quantified the scaling in distribution of selection intensities between, we can release our assumption of a simple scaling with the species-wide distribution and independently infer the distribution of effects for each DCA score bin and each sample from each cluster. The geometric mean of the deleterious effects is also strongly decreasing with DCA score when compared across pathotypes (Fig. 8B). Interestingly, even the pathotypes with the weakest selection efficiency show an increase in the mean deleterious effect with DCA score, highlighting a robustness of DCA predictions to varying population sizes. Note that as most mutations in the neutral bin are likely to be deleterious but with an effect too low to be counter-selected in these smaller populations, we computed their contribution to the deleterious mean as if their effect were in the range −1 ≤ 2*N*_*e*_*s* ≤ −0.1. A similar population-size dependent pattern can be observed on Codon scores when comparing a few clusters with thousands of isolates, covering different pathotypes, population size and inter-host transmission dynamics (Supplementary Fig. 5).

**Fig. 8:**
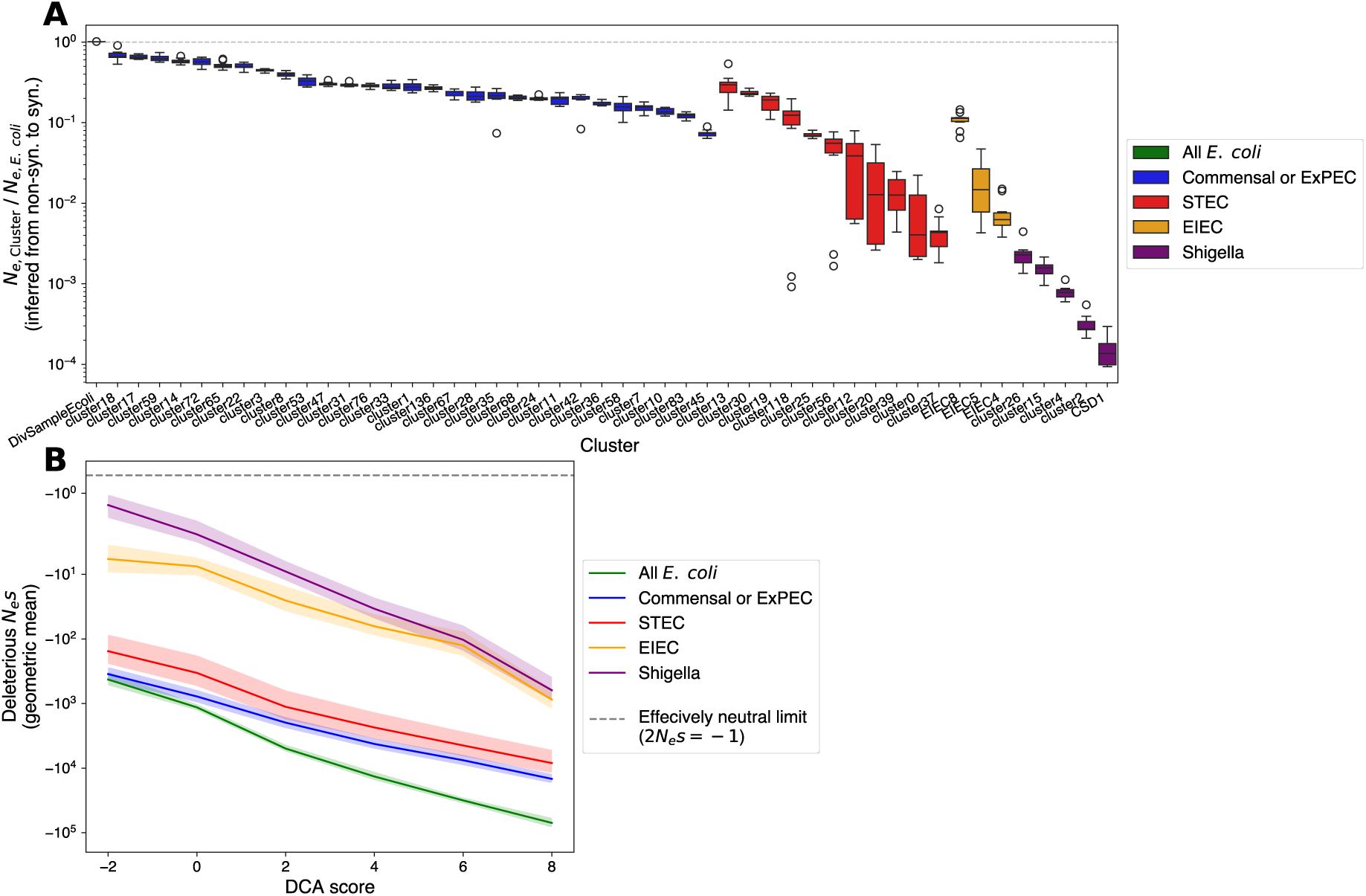
DFE inference for non-synonymous mutations in the main *E. coli* clusters. **A.** Selection efficiencies in various clusters as a fraction of the selection intensities inferred for the whole *E. coli* species, computed as the best scaling factor by while to multiply the *E. coli* DFE to best fit the cluster’s SFS, and interpreted as a population size ratio (*N*_*e*,Cluster_*s*⁄*N*_*e,E.coli*_*s*). **B.** Geometric mean of the deleterious selection intensities inferred for different *E. coli* lifestyles (averaged over all clusters with this lifestyle). 95% confidence interval is represented as a colored area.

## Discussion

The advent of widespread genomic sequencing has spurred the development of population genetics tools to infer key evolutionary processes from polymorphism data. In these methods, mutation frequency is a quantitative variable that can be directly estimated from genomic data, whereas selective effects are not directly observable and have to be estimated from polymorphisms frequency patterns such as the SFS. In parallel, advances in large-scale protein data analysis have driven the development of numerous bioinformatics tools that predict protein features, including the effects of protein variants.

In this study, we showed that large-scale genomic datasets such as our 81,440 *E. coli* genomes collection can serve as real-world benchmarks for evaluating mutation effect predictors, both for synonymous or non-synonymous mutations. This dataset had already been leveraged to demonstrate the improved predictive accuracy of DCA models compared to non-epistatic, independent-site protein models^20^. Our choice to keep using MSA-based DCA models instead of more recent generalist protein language models and other large machine learning approaches stems from the simplicity, flexibility and interpretability offered by DCA models. Indeed, DCA models are typically used to identify interactions and quantify mutation effects within a single protein domain or an entire protein, but this approach could be easily extended to capture any evolutionary constraints represented in the training MSA – even if it encompasses several interacting proteins^44^. Our approach is, to our knowledge, the first to apply population genetics as an analytical framework that exploits mutation frequency as a quantitative variable on such a large scale, rather than relying on a binary frequent/unfrequent classification defined by arbitrary thresholds most often used when assessing the predictive power of such bioinformatics tools.

Beyond assessing our capacity to predict mutational effects, our study also offers new perspectives on the mechanisms of natural selection at the species level. Most advanced population genetics methods have been developed for eukaryotic species, which typically feature low gene density and recombining linear chromosomes. These properties enable the distinction between neutral sites linked to selection in nearby genes and freely segregating neutral sites located further away. In contrast, bacterial genomes are highly compact with circular architecture, limited recombination constrained to double crossovers, and high codon usage bias. Quantifying selective effects along the bacterial chromosome is therefore key to deepening our understanding of long-term selective processes in prokaryotes.

Initially designed to infer the selective effect of a single class of mutations^40^, the PRF framework was soon extended to infer a full distribution of effect from a pair of SFS of non-synonymous and synonymous mutations^8^, as a single-effect model could not match the observed patterns. Considering a heterogeneous range of effects instead of a single one became a strength of this approach, as it can produce a result for any reasonable shape of synonymous and non-synonymous SFS. However it is also a weakness as it cannot test whether groups of mutations with similar effects behave as expected under the selection model used: it only describes what global distribution of effects would be compatible with the observed SFS. In other words, the inferred distribution of fitness effects is a direct mathematical transformation of the input data via a formula derived from the model, but doesn’t test whether we are right in using that model.

Our goal here was to categorize mutations using genetic scores so that mutations with similar effects could be regrouped together and fitted with the single-effect fitting framework. Our approach was successful for Codon scores, as we were able to estimate the effect of individual synonymous mutations in a deterministic way with their Codon score. As a comparison, the distribution-fitting methods can only infer the global shape of the distribution, but the effect of each individual mutations remains inaccessible. Our score-based approach however could not only determine the overall distribution of effects, but link it to a distribution of scores that can be explicitly computed for any synonymous mutation.

DCA scores, on the other hand, aim to capture selective constraints far more complex than codon preference, preventing us to use single-effect estimators for each DCA score value. However, both the mean effect and the proportion of mutations with mild effects of the distributions inferred with the distribution-fitting methods behave consistently with score values, with absolute mean effects increasing over several orders of magnitude as DCA score increases and the proportion of mild mutations and neutral plus beneficial mutations dropping to zero for mutations with DCA scores higher than 2 and 0, respectively. Rather than being interpreted as a direct proxy for individual non-synonymous effect, DCA scores are better understood as latent variables capturing the chances that an individual mutation be beneficial, neutral, mildly deleterious, or highly deleterious – with several nuances of high deleteriousness, that we can distinguish thanks to our large sample size. Still, they draw a connection between proteomics and population genetics by breaking down the abstract distribution of fitness effects into more concrete categories and making individual selective effects a bit more predictable. As predictors of selective effects continue to improve in accuracy, they may ultimately allow us to test foundational assumptions of population genetics models—such as the independence of segregating sites, constant selection coefficients, panmixia, or the limited role of epistasis – which are also predicted to impact the SFS^45^.

Finally, our large-scale quantitative framework helps clarify ongoing debates by providing meaningful orders of magnitude. Synonymous mutations are typically treated as a neutral reference in population genetic methods^10^, an assumption that is occasionally questioned by studies showing selection occurring on synonymous mutations^41,46^. Our results reconcile these perspectives by quantifying the distribution of codon-level effects. Consistent with previous estimates obtained from smaller datasets encompassing fewer genes, codons, and isolates^47^, we find that the majority of synonymous mutations have a small effect with *N*_*e*_*s* in the range [−1, 1] in the *E. coli* species, while non-synonymous mutations have selection intensities in the range [−10^6^, 1]. Codon selection results in a no more than 3-fold enrichment or depletion upon fixation, with the exception of the Leucine synonymous mutation CTA ↔ CTG (Codon score = ±2.9) showing higher effects. Moreover, this effect is discernible in the SFS of large and genetically diverse *E. coli* populations but becomes indistinguishable in smaller populations. This observation illustrates a central principle of population genetics: the impact of selection depends not only on the intrinsic effect of a mutation but also on the effective size of the population in which it segregates—a consideration often overlooked in experimental studies of synonymous mutations.

Likewise, our population genetics framework allows us to quantitatively assess the fundamental differences between InPEC such as *Shigella*, STEC/EHEC, and EIEC, and commensal/ExPEC lineages. InPEC strains are specialized pathogens that consistently cause gut dysbiosis, whereas ExPEC lineages consist largely of commensal gut residents capable of becoming opportunistic pathogens when translocated to extra-intestinal sites such as the urinary tract, the bloodstream or the meninges. Although both groups are predominantly represented by isolates obtained from pathogenic contexts, their population structures differ markedly. InPEC isolates derive mostly from low-diversity clades that have adapted to mammalian or human intestinal dysbiosis, while ExPEC isolates originate from far larger and more diverse populations occupying a wide range of environments, thereby reflecting the genetic diversity of their ancestral genome pools.

Our analysis quantitatively confirms the most extreme population size reduction of *Shigella* clusters previously described^28^ and extends it to other InPECs. We estimate the effective population sizes of *Shigella* clusters to range between one five-hundredth and one ten-thousandth of that of the full *E. coli* population. The lineage with the smallest inferred population size corresponds to *S. dysenteriae* serotype 1, presumably the most dangerous *E. coli*: it is a highly virulent strain that is both STEC and *Shigella*, and the original pathogen described by Kiyoshi Shiga in 1898^23^, explaining the overlapping nomenclature of these groups, both associated with the “Shiga” label due to either the presence of the Shiga toxin or the invasion plasmid. At the opposite end of the InPEC spectrum, the EIEC8/ST99 lineage from cluster 3 exhibits a much larger inferred effective population size—approximately one-fifth that of the entire *E. coli* species—and represents the most recent acquisition of the invasion plasmid within an *E. coli* background. This lineage, first identified in 2016^48^, still comprehends non-EIEC members with the same Sequence Type, while those that possess the invasion plasmid lack many of the secondary pathoadaptive mutations that characterize established *Shigella* and EIEC^49^.

Examining large genomic collections through the lens of population genetics enables quantitative estimates of key evolutionary processes. Beyond classical approaches such as the *Ka/Ks* ratio or the Poisson Random Field framework, complementary metrics from other disciplines—such as Direct Coupling Analysis (DCA) scores—can provide independent insights. Unlike diversity statistics, which depend on the number of observed mutations, or *Ka/Ks* ratios, which rely on their coding classification, DCA scores derive directly from the predicted impact of mutations on the proteins they encode. Across the different *E. coli* clusters, DCA enrichment slopes, *Ka/Ks* ratios, PRF-derived population sizes, and population diversity metrics show overall correlations, yet each captures distinct facets of selection (Fig. 9). *Ka/Ks* and PRF appear particularly sensitive for distinguishing differences in selection efficiency among low-diversity InPEC populations but tend to saturate in high-diversity commensal/ExPEC clusters. In contrast, DCA enrichment slopes are less correlated with diversity in InPEC clusters but show the strongest correlation with diversity among commensal/ExPEC clusters (*r*^2^ = 0.84, Supplementary Table S1). This pattern may reflect adaptive processes acting at different timescales: deleterious mutations at low frequency could represent short-term local adaptation before being purged over longer evolutionary intervals^17^. Consequently, DCA offers valuable potential for assessing the efficacy of selection across bacterial species—a scale at which *Ka/Ks* has limited resolution due to species-specific fitness landscapes and extensive genomic divergence^50^. Genomic predictors such as DCA and Codon scores therefore expand the set of tools available to quantify evolutionary dynamics, establishing a bridge between short-term, intra-specific population genetics and long-term processes of bacterial diversification and speciation.

**Fig. 9:**
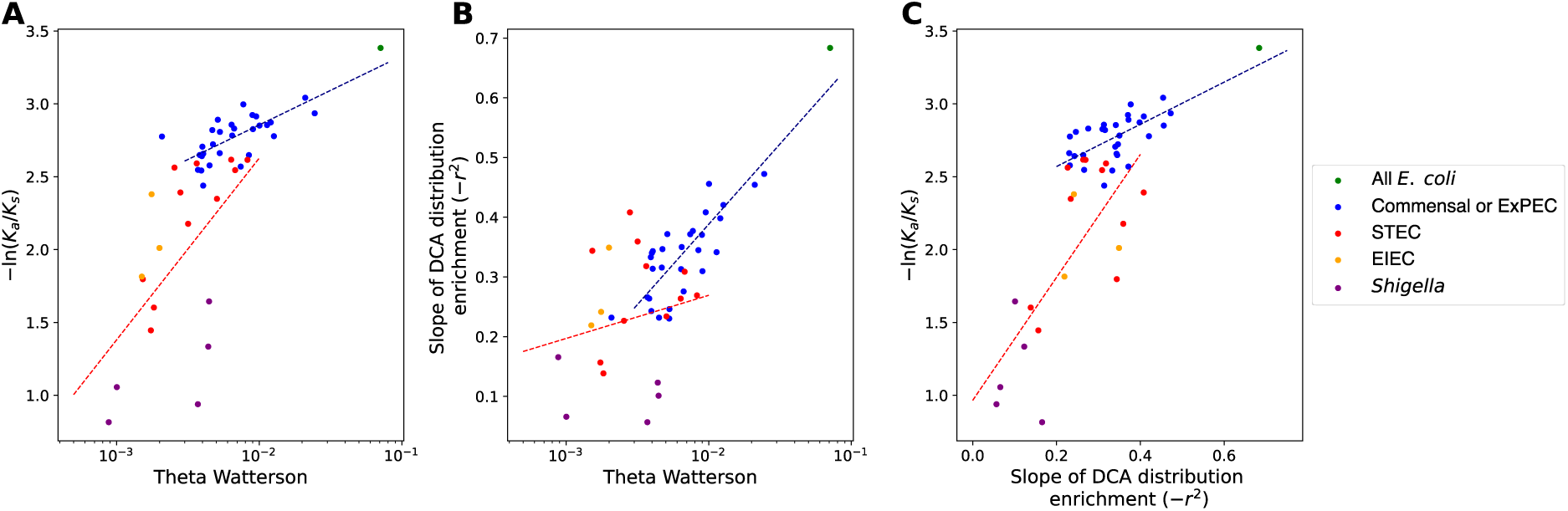
Comparison of correlations between selection intensity and diversity metrics. **A.** Linear regression between Ka/Ks on a log scale and the *θ*_W_diversity measurement. The regressions computed on the ExPEC/Commensal clusters, in blue, and the InPEC clusters, in red, are displayed. **B.** Linear regression between the DCA enrichment slopes from figure 4 and *θ*_W_, computed across the two same groups of clusters **C.** Linear regression between Ka/Ks and the DCA enrichment slopes computed across the two same groups of clusters.

## Methods

### 1. Dataset

A total of 82,063 contig-level genome assemblies were downloaded from the *Escherichia/Shigella* section of EnteroBase in February 2019^32,33^ and genomes corresponding to other *Escherichia* species and clades were excluded, leaving 81,440 *E. coli sensu stricto*^33^. Prodigal^51^ was used to identify 409,049,104 DNA open reading frames (ORFs) across the 81,440 genomes, leading to a mean 5,023 ORFs by genome, which is expected in *E. coli*. Quality filtering and several clustering steps using MMseqs2 and Mafft led to a pangenome of 349,553 protein sequences^17,52,53^. The consensus sequences of pan genes were then matched against the Swissprot database (October 2024 download). A Swissprot annotation was found for 12.1% of the proteins (42,412 out of 349,553), but this figure increased significantly when considering genes present in the majority of genomes. All extracted gene sequences and their translation, genomic location and annotation were stored in a SQL database using Python3.8 with the sqlite3 and pandas libraries.

### 2. Core genome, genome clustering and phylogenetic reconstruction

A core genome alignment (>95% presence, <1% duplicate) of 3,016 genes was used to create a genetic distance matrix, measured as the proportion of non-gapped bases in the alignment. Note that genomes harboring the *Shigella* marker ipaH3 were excluded for the selection of core genes. The resulting distance matrix (6.6 billion coefficients, 36 Gb) has a mean of 1.66% (on non-diagonal coefficients) and a maximum of 2.97%. We clustered it with the DBSCAN algorithm^54^ with parameters: epsilon=0.5% and minimum samples=5 and obtained 240 clusters of variable size, ranging from 5 genomes to 11,768, that together contain 80,843 genomes (99.2% of the database). The 597 remaining genomes could be further regrouped into 339 additional, uncommonly sampled clusters containing 1 to 4 genomes. We also used this matrix to create a ‘diversity sample’ of *E. coli*, consisting of 718 genomes that are distant by more than 0.475% from each other (95% of the 0.5% cluster distance), which is more representative of the species *E. coli* diversity as genomes sampled randomly from the complete database would be biased towards pathogenic lineages. For each cluster with more than 110 genomes, 100 genomes were randomly drawn and polymorphic sites present in each one of these subsamples were detected and oriented similarly as in the whole *E. coli* species (see section 4). This process was repeated 10 times for each cluster, defining

As building a phylogeny of the entire database was unconceivable, we used the following procedure: for each one of the 240 main clusters, a phylogeny was built from the core genome with FastTree^55^ and ancestral state reconstruction (ASR) with IQTree^56^ allowed to derive the most likely ancestral core genome. A phylogeny of the 240 resulting reconstructed core genomes was then built using IQTree. Gubbins^57^ was run prior to all phylogenetic inference to detect and mask recombined fragments.

### 3. *E. coli* classification, pathotype determination and additional genomes

ST and ST complexes were determined by using the *mlst* software^58,59^ and by downloading the Pubmlst ST Complex reference^59^ (as of 02/05/2025). Phylogroups had been previously determined^33^ using both the ClermontTyping^60^ and the Mash^61^ genome-clustering methods. Serotypes were determined with ECtyper^62^.

*Shigella* had been previously detected using the presence of the ipaH3 marker^33^, present on the pINV plasmid and sometimes translocated in the *Shigella* and EIEC chromosome^63^. In order to obtain a more precise and up-to-date annotation, we applied ShigEIFinder^27^ to the 81,440 genomes of the database. This software identifies and regroups 59 *Shigella* serotypes and 22 EIEC serotypes into a few large genetic clusters: the *Shigella sonnei* cluster and 3 large clusters containing 54 of the 58 remaining *Shigella* serotypes. Named C1, C2 and C3, we refer to them as *Shigella 1*, *Shigella* 2 and *Shigella 3* in order not to create confusion with our own genomic clusters. A summary of the classification according to serotype, ST and *Shigella*/EIEC clusters is available for each cluster in Supplementary Table 2.

Using the ShigEIFinder annotation, we identified several clusters or groups of related genomes within a cluster annotated as *Shigella* or EIEC that did not reach the required quorum of 110 isolates: *S. dysenteriae 1*, *S. dysenteriae 8*, *S. boydii 12*, and 5 EIEC linages called C4-C8 by ShigEIFinder and that we rename EIEC4-EIEC8. We chose to enrich our database for these particular lineages from the current (June 2025) version of Enterobase, which has expanded from 82,063 to more than 380,000 genomes in 6 years. Using the ST corresponding to each ShigEIFinder annotation, we were able to download more than 100 genomes for *S. dysenteriae 1* and for 3 EIEC lineages: EIEC4 / ST270-ST311 (cluster 41), EIEC5 / ST6 (part of cluster 8), and EIEC8 / ST99 (part of cluster 3). As these last two represent less than 1% of their parent cluster, we treated them as independent populations in the rest of the analysis.

In order to detect STEC and EHEC clusters, we used the Swissprot annotation to detect genes annotated as Shiga toxin coding-genes (*stxA*, *stxB*, *stxA2* and *stxB2*), hemolysins (*ehxA*), and intimins (*eae*). We then classified clusters which contained a *stx* gene with more than 50% prevalence as STEC, and clusters that also contained a majority of genomes with *ehxA* and *eae* as EHEC. We found that the largest of these clusters correspond to serotypes that are indeed described as STEC and EHEC in the litterature^64^. For the statistical analysis, we regrouped all STEC and EHEC in a single category STEC-EHEC.

### 4. Training DCA models, detecting and orienting mutations

In order to compare distinct *E. coli* clusters including *Shigella*, we selected a core genome of the entire *E. coli* species, resulting in 2,652 genes. We attempted to find a homologous sequence in the outgroup species *Escherichia albertii* (GenBank CP025317), *Escherichia fergusonii* (GenBank CU928158) and *Salmonella enterica* (Mage VFAF01.1) and train a DCA model for each of the genes that was shorter than 800 amino-acids. DCA models were trained precedently^17^ by retrieving a Multiple Sequence Alignment (MSA) with HHblits^65^ and then using the Julia library plmDCA v.0.4.1^66^ to train the model. Homologous sequences with more than 90% identity with their corresponding *E. coli* gene were removed, only genes for which the MSA contained more than 200 homologous sequences were kept, and we removed sites for which less than 100 homologous sequences presented a gap at this position. We retained 2,358 genes with a DCA model and at least one outgroup sequence, but the majority of them (92.6%) actually possess two to three outgroups.

We generated all possible single-amino-acid variant at each site for each protein and measured its DCA score difference with the score of the consensus. This set of mutation scores was then weighted by the probability of such an amino-acid substitution occurring via a single base substitution given the *E. coli* codon usage and its base substitution rates (Supplementary Fig. 6). In order to estimate these base substitution rates prior to any selection, we counted the number of singleton mutations for each possible base substitutions in the ‘Diversity Sample’ dataset. Using low frequency (1 in 718) mutations allows to estimate the mutation biases before selection had the time to significantly distort them, and using a set of diverse *E. coli* allows to capture the species-wide mutation biases, that could fluctuate in some lineages. Codon usage was derived by determining the most prevalent codon among all alleles of each gene and counting them. By coupling codon usage with mutation bias, we could determine a 30.3%-69.7% synonymous / non-synonymous mutation proportion. Without more knowledge about the selective pressure on each core gene, we aggregated all mutations in a single weighted distribution of the DCA scores arising by random mutation, prior to any selection.

For the Codon scores, such an approach was impracticable as the scores depend only on the codon substitution, without a position and context-dependency layer, meaning that small fluctuations in mutation biases across clusters or reading frames will have large impact on the distribution (Supplementary Fig. 7 and 8). Thus, for each dataset, we computed the ‘local’ distribution of Codon scores as the distribution of scores at low frequency (*n*_*mut*_ ≤ 3 for the ‘Diversity Sample’ and 3 ≤ *n*_*mut*_ ≤ 5 in for the entire database in Fig. 5 and 6, and proportion of mutations in the first frequency bin for Supplementary Fig. 5).

### 5. DCA scores and Codon scores

DCA is currently employed as a protein mutational landscape model: it predicts the functionality of any amino-acid sequence relative to that of the homologous protein family it was trained on. This functionality is quantified as a statistical energy *E*(*a*_1_, *a*_2_, … *a*_*L*_) for an amino-acid sequence *a*_1_*a*_2_ … *a*_*L*_ of length *L*, which is linked to the probability of observing a variant of the protein with this sequence by the formula: 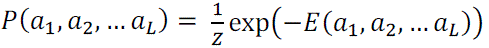 where *Z* is a normalization constant taken over all possible sequences of length *L*. When the model is trained correctly on a set of homologous proteins of the same family, non-functional sequences and sequences that don’t belong to this protein family are attributed very low probability and very high energy *E*(*a*_1_, *a*_2_, … *a*_*L*_), while protein from the training set and new proteins of the same family are attributed higher probabilities and lower energies. This statistical energy is the sum of a contribution from all amino acids at making the sequence more or less protein family-like, and a contribution from all pairs of amino acids across sites. More formally, a DCA model consists in a site-dependent bias matrix *h* and a pairwise coupling matrix *J* such that:

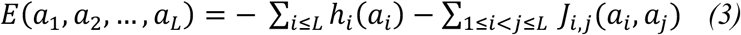

Although energy is defined for an entire sequence, we can easily compute the change in statistical energy associated with a single mutation from amino acid 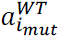 to amino acid 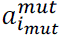 at site *i_mut_* given the amino acids at the other sites, by taking the differences between the energy of the original sequence and the energy of the same sequence with one substitution at site *i_mut_*. This energy difference is called the DCA score of the mutation:

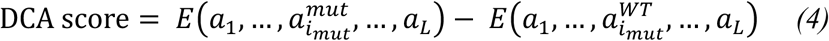

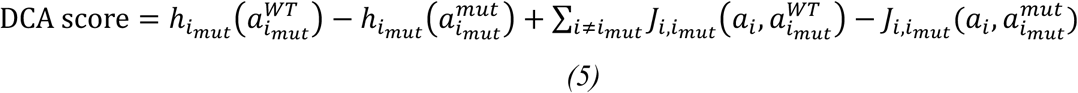

The first difference captures the importance of a specific amino-acid at site *i_mut_*, for instance due to polarity constraints in transmembrane proteins or enzymatic sites specificities, while the second term captures the interactions between the focal site and the rest of the sequence and can account for protein-wide epistasis. Although with this definition a DCA score is always defined in a specific context – *i.e.* for a given set of amino acids at the other positions – in practice a few substitutions at other sites will not dramatically affect the value of the DCA score of a substitution the focal position: context-dependency of the score value of an individual substitution gradually builds-up with sequence divergence and is very low between sequences with less than 5% amino-acid variation such as *E. coli* genes^20^. This allows us to speak about “the” DCA score of an amino-acid substitution in “the” *E. coli* context, which justifies our approach of computing all substitution scores within the context of the *E. coli* consensus allele for each gene.

Codon scores were defined as explained in the Results by counting the number of each codon in a concatenation of the consensus DNA sequence of the 2,358 selected core genes, and for each possible synonymous mutations we computed the score as the opposite of the logarithm between mutated and ancestral codon usage, so that negative score reflect better-suited codons, as for DCA scores.

### 6. Choice of a selection model and its interpretation

We use a simple mathematical model accounting for both selection and drift of recurrent mutations segregating independently in a panmictic population, known as the Wright-Fisher model. In this model, each mutation possesses a selective effect 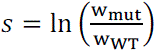 were w_WT_ is the Wrightian fitness (expectation of the number of offspring) of a wild-type genotype and w_mut_ the fitness of a genotype carrying a mutation (here representing a single codon substitution) but with an otherwise identical genome. The selective effect is supposed to be constant (environment-independent) and independent of the presence of subsequent mutations. Note that this is simply a modeling approximation and does not mean that the model is incompatible with the effects of many mutations being environment-dependent nor with epistasis being widespread in the *E. coli* genome. The constant selective effect *s* captured by our model can rather be interpreted as the selective effects of a mutation in different environments averaged by the proportion of time a typical *E. coli* lineage spends in each environment, and similarly if the effect changes when it is associated with different subsequent mutations at other sites, what we capture corresponds to an average over frequently observed subsequent mutations.

The population is characterized by its equivalent population *N*_*e*_ that represents the number of individuals effectively contributing to the following generations in the long run and can be several orders of magnitudes lower than the actual number of individuals at a given time or census size *N* which rather represents the ecology of the species. For instance, estimates for the *E. coli* census size is of the order of *N*∼10^20^ ^67^ but this population is highly structured: the gut of a mammalian host typically harbor a few strains of millions to billions individuals sharing the *E. coli* niche and there are hundreds of billions to trillions of such mammalian hosts, with occasional migration from one host to another as well as less documented but equally important environmental niches such as water treatment plants and waterways. The chances of any particular *E. coli* lineage being successively translocated to and adapting to a series of new niches are thus relatively small, explaining why population geneticists have observed a genetic diversity corresponding to what would be observed in a panmictic population of equivalent size *N*_*e*_∼10^8^ to 10^9^.

### 7. Inference of selection intensity with the Wright-Fisher model

With these nuances in head, we use the diffusion approximation of the Wright-Fisher model and its prediction of the equilibrium distribution of mutations frequency *q* ∈ [0,1] (continuous version of the site-frequency spectrum) of recurrent mutations with selective effect *s* ^68–70^:

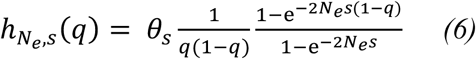

represents the expected number of dimorphic sites with selective effect *s* of the mutant allele found at frequency *q*. Here, *θ*_*s*_ = 2 *N*_*e*_*μ*_*s*_ = 2 *N*_*e*_*μ ρ*(*s*) corresponds to twice the rate of apparition in the population of new mutations with effect *s* by genomic site (*μ* is the mutation rate by individual by genomic site and *ρ*(*s*) is the distribution of new selective effects (DFE), corresponding to the probability that a new mutation appears with effect *s*).

When genetic polymorphism sequencing data became available in the 1990s, they were reduced to a few genes sequenced in a few individuals, motivating the development of a probabilistic framework able to deal with the uncertainty arising from these relatively small sample as well as adapting a continuous equation to discrete data. This framework is the Poisson Random Field (PRF) approach: given a sample of size *n* individuals, the number of mutated sites found in *i* individuals can be modelled as a Poisson variable with an expectation *F*(*i*, *n*, *θ*_*s*_, *γ*_*s*_) depending on the sampling size *n*, the absolute frequency *i*, the selection intensity

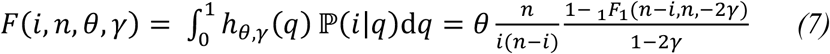

where _1_*F*_1_ is the confluent hypergeometric function.

The values of *θ* and *γ* could then be inferred from the observed SFS by maximum likelihood^40^. However, the SFS can also be affected and distorted by demographic processes, such as population size variations and population structure^34^, that could bias selection inference if not handled correctly. As these processes will impact the SFS of all mutations currently segregating, contrasting the SFS of different kinds of mutations with the PRF is a solution that has been successfully applied at estimating the distribution of selective effects of non-synonymous mutations, using non-selected mutations to correct for distortions in the SFS^8^. This procedure requires to have good estimates of the proportions of new selected and non-selected polymorphisms arising by mutation.

In our PRF approach, we defined the expected SFS ratio of selected over reference neutral mutations:

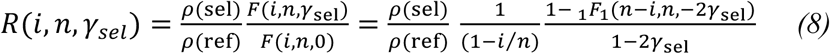

When fitting a single selective effect such as with Codon scores for synonymous mutations, we minimized the following log-likelihood function, with *k*_sel_(*i*) and *k*_ref_(*i*) denoting the SFS of the selected and the reference mutations, respectively:

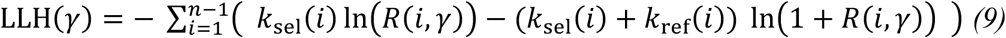

The terms within the sum correspond to the likelihood of sampling *k*_sel_(*i*) selective mutations from a binomial law ℬ(*n* = *k*_sel_(*i*) + *k*_ref_(*i*), *p*) when *p*⁄(1 − *p*) = *R*(*i*, *γ*).

When fitting a distribution of effects with several bins, the parameters to minimize were the log-probabilities of each bin. The bins were defined as powers of 10 of 2*N*_*e*_*s* except around 0: 2*N*_*e*_*s* ∈ [−10^6^, −10^5^]; [−10^5^, 10^4^]; …; [−10, −1]; [−1,1]; [1,10] and the effects were uniformly distributed within each bin on a log scale except for the effectively neutral bin in which they were uniformly distributed on a linear scale. The log-likelihood sum was then taken both on *i* and on the bins (weighted by their proportion in the DFE of the selected mutation). All bins with deleterious selective effects stronger than 10 × *n*_sample_ were merged in a single bin (for the *n* = 718 ‘Diversity Sample’ and the *n* = 100 cluster samples).

The curve fit method was very similar in its spirit but relying on fitting the following function with log-Huber loss:

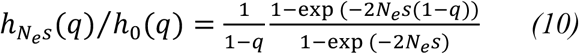

This is the ratio of the selective to neutral equilibrium distributions of mutations frequencies. The frequencies were discretized on a logit bin as are the points in Fig. 6A and 7B, and the quantities to fit corresponded to the ratio of the number of observed selected to reference mutations observed within each bin, divided by the ratio of the probability of apparition of these mutations.

We took synonymous mutations with a Codon score in [−1,1] as reference mutations in Fig. 6 and 7 and all synonymous mutations in Fig. 8, due to the lower total number of mutations in the small clusters.

### 8. Implementation and data treatment

Data were treated in Python v3.9 using the BioPython, numpy and pandas libraries, and figures were realized using the matplotlib, seaborn, statannotations, and plotly libraries and the Inkscape software. We used scipy v1.14.1 for the selection intensities inference. Data treatment and storage were conducted on the French Bioinformatics Institute’s platform (IFB Core Cluster).

## Data Availability

The datasets were derived from sources in the public domain: Enterobase (https://enterobase.warwick.ac.uk/), Uniprot30, MAGE and GenBank. Intermediate data files comprising MSA and polymorphism datasets will be made available on Zenodo.

## Code Availability

All codes to extract genetic sequences, build SQL database and analyze polymorphism will be made available on GitHub.

## Acknowledgements

Our work was partially funded by the French Agence Nationale de la Recherche EcoRecEp grant (ANR-23-CE35-0006, to O.T.)n DREAM grant (20-PAMR-0002, to O.T.) and PROTEVOL grant, the French Fondation pour la Recherche Médicale (SMC202505021060), and the PhD program AMX of École Polytechnique and Ministère de l’Enseignement Supérieur, de la Recherche et de l’Innovation (to M.M.). All authors are grateful to the French Bioinformatics Institute (IFB) for housing the data and providing computational tools and support.

## Author Contributions Statement

O.T. and M.W. designed the analysis. M.M. and L.V. did the bioinformatics treatment and contributed to the analysis. G.C. contributed to the analysis. M.M. and O.T. wrote the paper.

## Supplementary Figures Legends

**Supplementary Fig. 1: Polymorphic sites used in the study. A.** Number of polymorphic sites within the 81,440 genomes database (at least two different codons, gaps and invalid codons removed), core polymorphic sites (one valid codon present in at least 95% of the genomes), nearly dimorphic sites (two major codons present in at least 95% of the genomes), and oriented polymorphic sites (one and only one of the two major codons is equal to one of the outgroup codons). The last bar represents the proportion of synonymous vs non-synonymous polymorphisms within the oriented polymorphisms. **B.** Same statistics as a proportion of Total polymorphism across the 49 *E. coli* subpopulations studied (45 main clusters + 4 additional EIEC and *Shigella* clades + ‘DivSampleEcoli’). The vertical bars represent the dispersion between these subpopulations.

**Supplementary Fig. 2: Histogram of the pairwise genetic distances within the 81,440 *E. coli* genomes.** The threshold of 0.5% was chosen as a neighborhood threshold for the genome clustering step.

**Supplementary Fig. 3: Summary of the STEC/EHEC clusters present in the database.** STEC (Shiga-toxin producing *E. coli*) are defined by the presence of the genes *stxA*-*stxB* (Shiga toxin) and/or *stxA2*-*stxB2* (Shiga-like toxin). The medical community is particularly concerned with a subtype of STEC called EHEC (entero-hemorrhagic *E. coli*) that is characterized by the production of a toxin called hemolysin (*ehxA* gene), and that also often encodes an attachment and effacement (*eae* gene) protein called intimin that allow the bacteria to stick closely to enterocytes. We report here the prevalence of these virulence factors in the clusters we defined as STEC/EHEC (*ie* those in which a majority of isolates possess a *stx* gene), as well as their serotypes.

**Supplementary Fig. 4: Theoretical SFS and SFS ratio with neutral reference for different values of selective effects.** The curves are representations of the formula 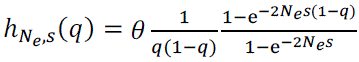 for the SFS, with *θ* = 0.08 as in the *E. coli* species, and 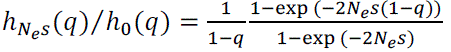 for the SFS ratio. We use a logit scale on the x-axis and a log scale on the y-axis for a better visual appreciation of the characteristic values of each curve (frequency of deviation from neutrality, final enrichment / depletion).

**Supplementary Fig. 5: Site frequency spectra for DCA scores and Codon scores of 4 clusters from the database:** cluster 0 (**A** and **E**) containing the most prevalent STEC/EHEC (ST11 complex – serotype O157:H7 – from phylogroup E); cluster 1 (**B** and **F**) containing the the most prevalent ExPEC (ST131 – from phylogroup B2); cluster 2 (**C** and **G**) containing the most prevalent *Shigella* (*S. sonnei* – within phylogroup B1 in the phylogeny); and cluster 3 (**D** and **H**) which is a much more diverse cluster containing a majority of genomes unassociated with any pathogeny (as well as a minority of STEC/EHEC O104:H21, O104:H4 and O146:H21, and a very small proportion of EIEC ST99). More information on the clusters is available in Supplementary Table 2.

**Supplementary Fig. 6: Mutation rates biases observed by counting the number of oriented base substitutions at low frequency** (3 to 5 genomes within the 81,440 of the database). The rates are scaled by the mean rate across all possible base substitution.

**Supplementary Fig. 7: Mutation rates biases observed across a few *E. coli* clusters.** The rates are scaled by the mean rate within each population, and the error bar represents the fluctuations between the 3 reading phases.

**Supplementary Fig. 8: Density of DCA scores of random mutations and newly arising mutations.** In black, the distribution of scores of all possible amino-acid substitutions (deletions excluded). In purple, the distribution of scores of amino-acid substitutions achievable with a single base substitution. In blue, the same distribution but where each substitution is weighted by the base mutation rate, taking into account the codon composition for each amino-acid and the base substitution rates. In light blue, the distribution of scores observed at low frequency (3 to 5 genomes in the 81,440 database).

**Supplementary Fig. 9: Ratio of the regression slopes presented in Figure 9 by the regression slope computed across all clusters** (whatever their pathotype), showing which parts of the clusters drive the regression for DCA enrichment slopes and for *Ka/Ks*.

## Supplementary Tables

**Supplementary Table S1: Correlations between diversity and selection efficiency measures over different groups of populations.**

**Supplementary Table S2: Summary statistics and description of the 579 genomic clusters in the 81,440 genomes database.**

