## Supplementary figures, tables S1 and S2 for "Linking Codon- and Protein-Level Mutation Scores to Population Genetics Reveals Heterogeneous Selection Efficiency Across *Escherichia coli* Lineages": Supplementary Figures.pdf

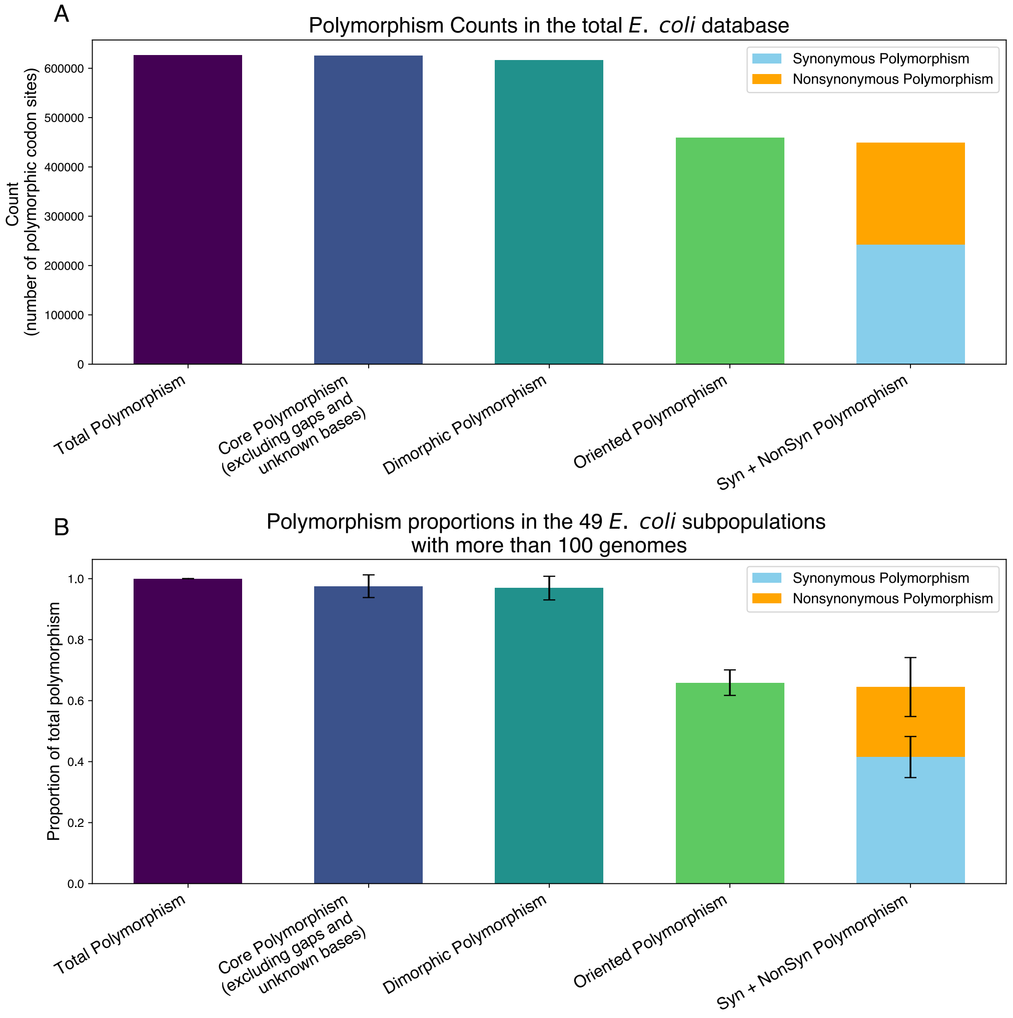


***Supplementary Fig. 1: Polymorphic sites used in the study. A.*** *Number of polymorphic sites within the 81,440 genomes database (at least two different codons, gaps and invalid codons removed), core polymorphic sites (one valid codon present in at least 95% of the genomes), nearly dimorphic sites (two major codons present in at least 95% of the genomes), and oriented polymorphic sites (one and only one of the two major codons is equal to one of the outgroup codons). The last bar represents the proportion of synonymous vs non-synonymous polymorphisms within the oriented polymorphisms.* ***B.*** *Same statistics as a proportion of Total polymorphism across the 49 E. coli subpopulations studied (45 main clusters + 4 additional EIEC and Shigella clades + DivSampleEcoli). The vertical bars represent the dispersion between these subpopulations.*

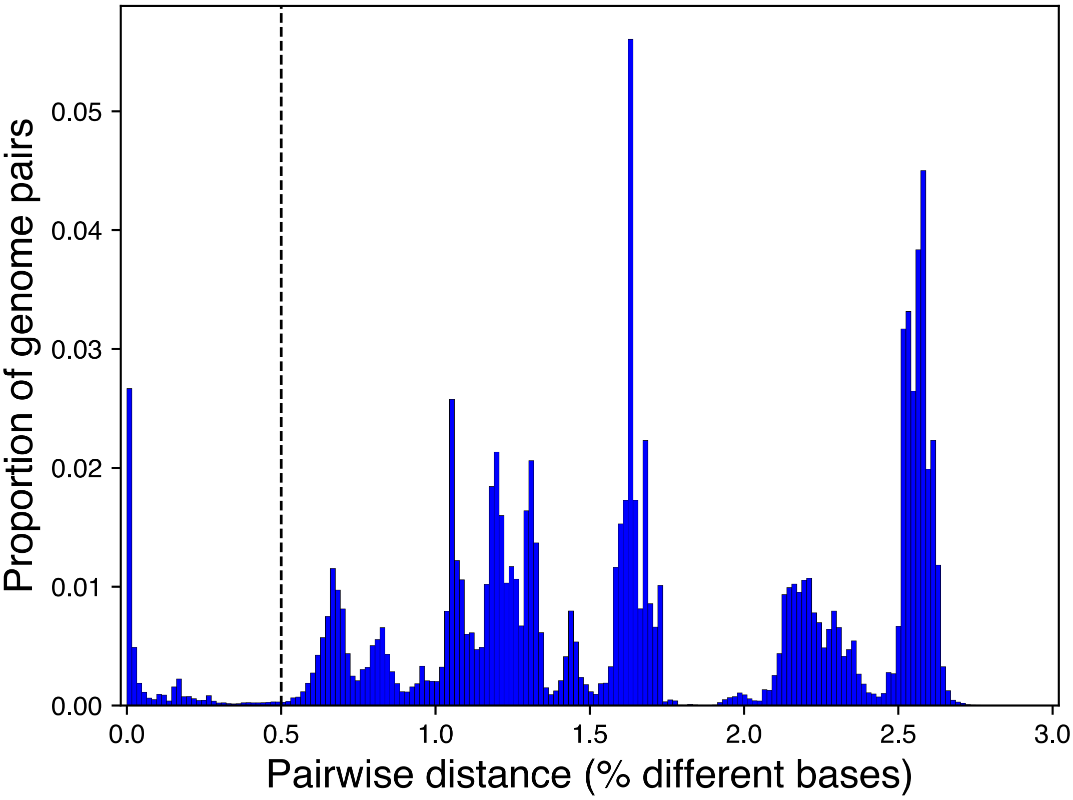


***Supplementary Fig. 2: Histogram of the pairwise genetic distances within the 81,440 E. coli genomes.*** *The threshold of 0.5% was chosen as a neighborhood threshold for the genome clustering step.*

| Cluster | Phylogroup | Sequence Types | Serotypes | Main 7 EHEC serotypes | Shiga toxin | Attachment and effacement | Hemolysin | Isolates_in_Database | Average pairwise distance | Maximal pairwise distance |
|---|---|---|---|---|---|---|---|---|---|---|
| 0 | E | ST11 (95.7%), ST335 (1.1%) and 71 other STs | O157:H7(90.7%), -:H7(8.1%), O55:H7(1.1%), O157:-(0.0%), and 2 other serotypes | O157:H7(90.7%) | stxA2 (67.5%), stxB2 (94.9%) | Intimin (99.5%) | ehxA (91.4%) | 10430 | 0.016% | 0.193% |
| 12 | B1 | ST21 (58.9%), ST16 (26.8%) and 101 other STs | O26:H11(46.8%), O111:H8(25.8%), -:H11(8.6%), -:H8(4.5%), and 39 other serotypes | O26:H11(46.8%), O111:H8(25.8%) | stxA (90.9%), stxB (91.1%) | Eae protein (62.4%), Intimin (67.5%) | ehxA (87.2%) | 6195 | 0.081% | 0.257% |
| 13 | B1 | ST17 (76.2%), ST1967 (10.7%) and 76 other STs | O103:H2(60.0%), -:H2(10.3%), O45:H2(8.4%), O123/O186:H2(6.6%), and 47 other serotypes | O103:H2(60.0%), O45:H2(8.4%) | stxA (88.7%), stxB (88.7%) | Intimin (89.3%) | ehxA (80.6%) | 3667 | 0.063% | 0.677% |
| 19 | B1 | ST655 (90.4%), ST2952 (3.9%) and 19 other STs | O121:H19(92.8%), -:H19(3.8%), O38:H26(0.8%), -:H9(0.6%), and 12 other serotypes | O121:H19(92.8%) | stxA2 (90.3%), stxB2 (90.4%) | Intimin (93.3%) | ehxA (84.0%) | 972 | 0.057% | 0.672% |
| 20 | E | ST32 (90.7%), ST137 (6.7%) and 5 other STs | O145:-(68.6%), -:-(31.2%), O117/O107:-(0.2%) | Probably O145:H28 (ST32 matching but H28 not recognized by Ectyper) | stxA2 (69.2%), stxB2 (70.8%) | Intimin (99.7%) | ehxA (89.6%) | 624 | 0.016% | 0.153% |
| 25 | B1 | ST342 (49.6%), ST119 (21.4%) and 21 other STs | O5:H9(43.4%), O165:H25(19.7%), O177:H25(13.8%), O145:H25(7.4%), and 9 other serotypes | Not found | stxA (58.5%), stxB (58.7%), stxB2 (52.4%) | Intimin (68.6%) | ehxA (82.8%) | 458 | 0.282% | 0.617% |
| 30 | B2 | ST583 (55.9%), ST722 (11.3%) and 14 other STs | O63:H6(38.2%), O145:H34(19.7%), O125:H6(16.4%), O132:H34(9.7%), and 8 other serotypes | Not found | stxA2 (62.6%), stxB2 (62.6%) | Intimin (83.6%) | Not found | 238 | 0.353% | 0.807% |

***Supplementary Fig. 3: Summary of the STEC/EHEC clusters present in the database.*** *STEC (Shiga-toxin producing E. coli) are defined by the presence of the genes stxA-stxB (Shiga toxin) and/or stxA2-stxB2 (Shiga-like toxin). The medical community is particularly concerned with a subtype of STEC called EHEC (entero-hemorrhagic E. coli) that is characterized by the production of a toxin called hemolysin (ehxA gene), and that also often encodes an attachment and effacement (eae gene) protein called intimin that allow the bacteria to stick closely to enterocytes. We report here the prevalence of these virulence factors in the clusters we defined as STEC/EHEC (ie those in which a majority of isolates possess a stx gene), as well as their serotypes.*

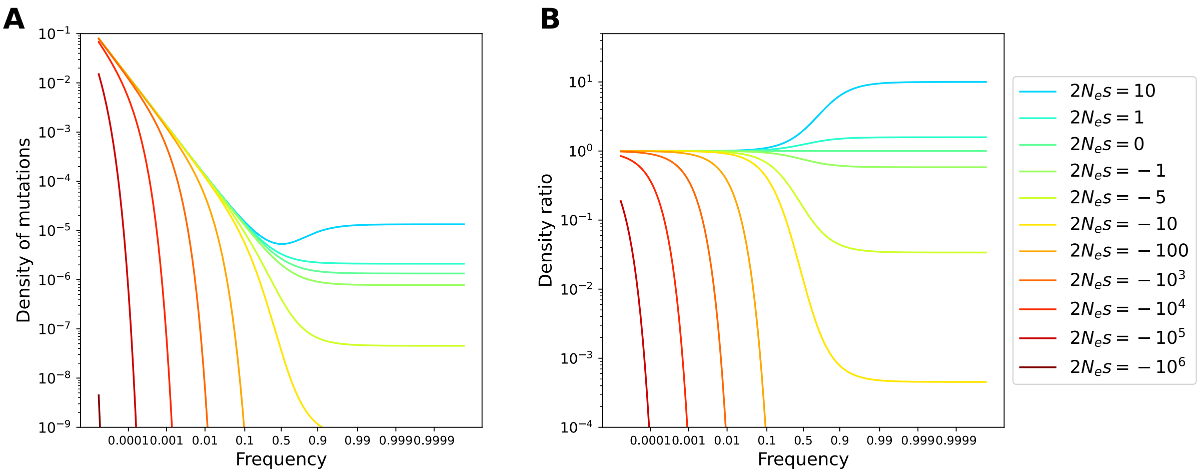


***Supplementary Fig. 4: Theoretical SFS and SFS ratio with neutral reference for different values of selective effects.*** *The curves are representations of the formula* $h_{N_e,s}(q) = \theta \frac{1}{q(1-q)} \frac{1-e^{-2N_es(1-q)}}{1-e^{-2N_es}}$ *for the SFS with* $\theta = 0.08$ *as in the E. coli species, and* $h_{N_es}(q)/h_0(q) = \frac{1}{1-q} \frac{1-exp\,(-2N_es(1-q))}{1-exp\,(-2N_es)}$ *for the SFS ratio. We use a logit scale on the x-axis and a log scale on the y-axis for a better visual appreciation of the characteristic values of each curve (frequency of deviation from neutrality, final enrichment / depletion).*

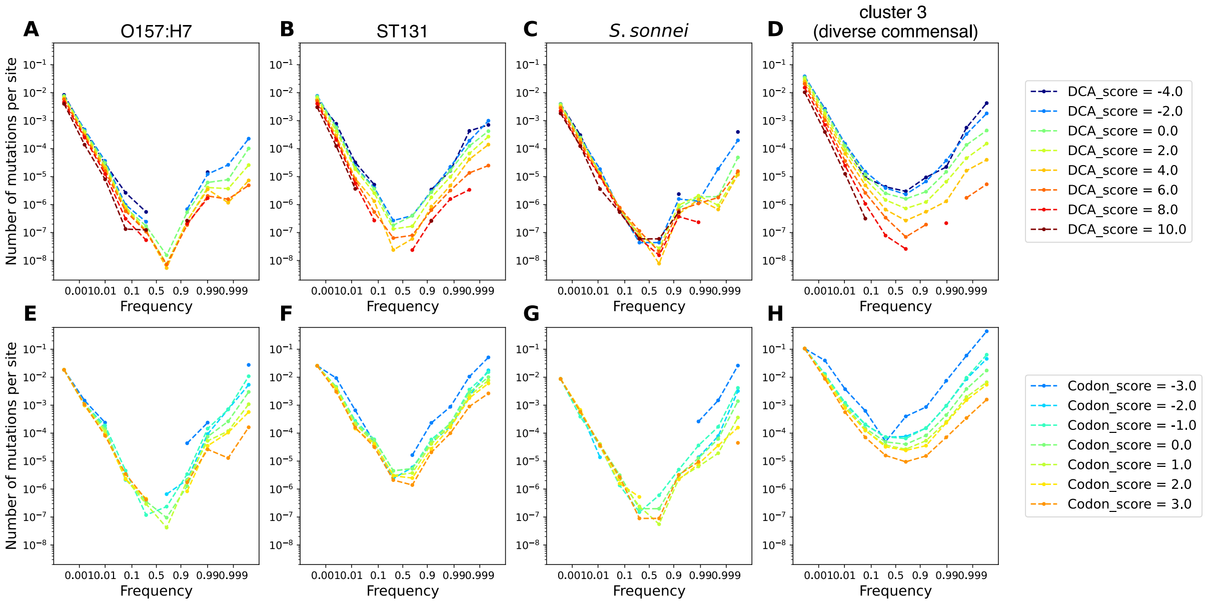


***Supplementary Fig. 5: Site frequency spectra for DCA scores and Codon scores of 4 clusters from the database****: cluster 0 (**A** and **E**) containing the most prevalent STEC/EHEC (ST11 complex – serotype O157:H7 – from phylogroup E) ; cluster 1 (**B** and **F**) containing the the most prevalent ExPEC (ST131 – from phylogroup B2) ; cluster 2 (**C** and **G**) containing the most prevalent Shigella (S. sonnei – within phylogroup B1 in the phylogeny) ; and cluster 3 (**D** and **H**) which is a much more diverse cluster containing a majority of genomes unassociated with any pathogeny (as well as a minority of STEC/EHEC O104:H21, O104:H4 and O146:H21, and a very small proportion of EIEC ST99). More information on the clusters is available in Supplementary Table 2.*

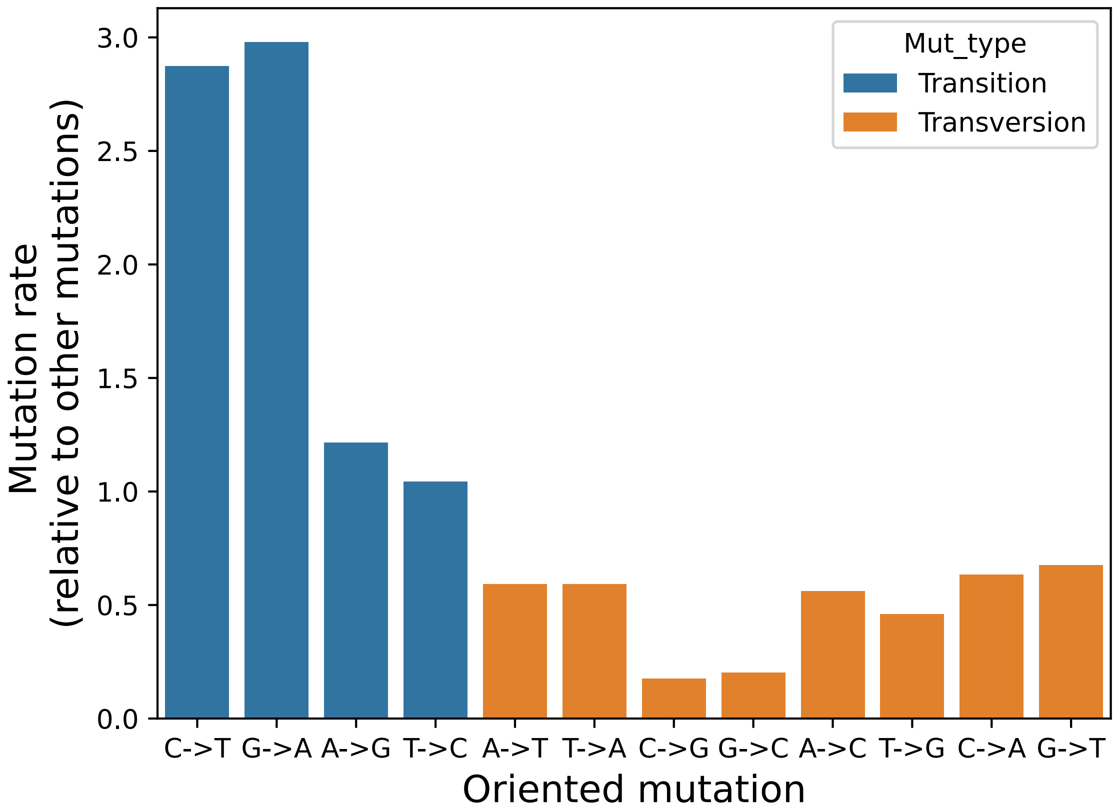


***Supplementary Fig. 6: Mutation rates biases observed by counting the number of oriented base substitutions at low frequency*** *(3 to 5 genomes within the 81,440 of the database). The rates are scaled by the mean rate across all possible base substitution.*

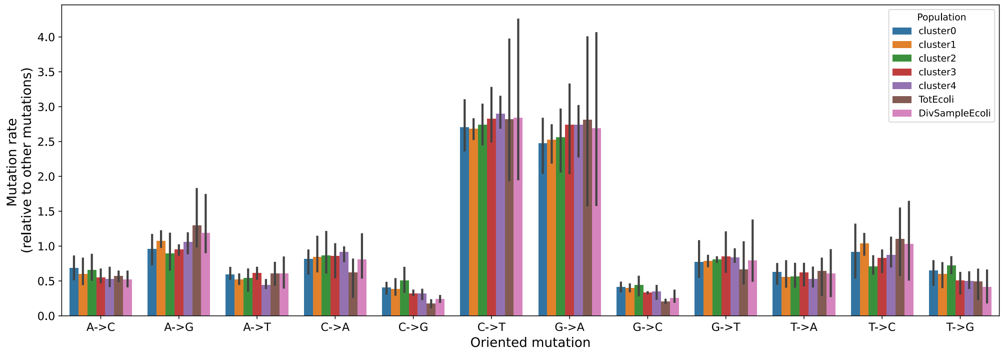


***Supplementary Fig. 7: Mutation rates biases observed by across a few E. coli clusters.*** *The rates are scaled by the mean rate within each population, and the error bar represents the fluctuations between the 3 reading phases.*

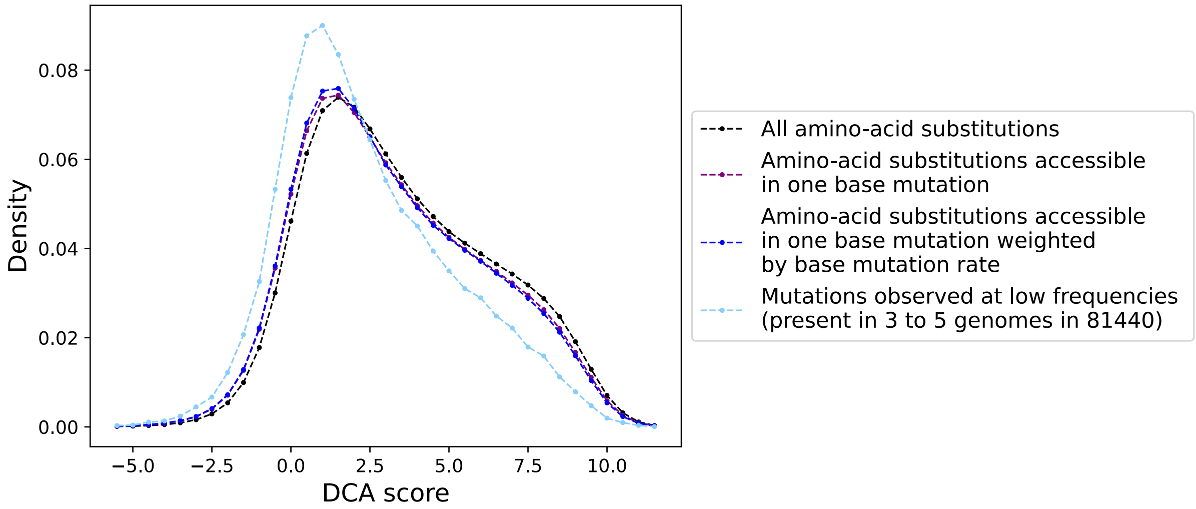


***Supplementary Fig. 8: Density of DCA scores of random mutations and newly arising mutations.*** *In black, the distribution of scores of all possible amino-acid substitutions (deletions excluded). In purple, the distribution of scores of amino-acid substitutions achievable with a single base substitution. In blue, the same distribution but where each substitution is weighted by the base mutation rate, taking into account the codon composition for each amino-acid and the base substitution rates. In light blue, the distribution of scores observed at low frequency (3 to 5 genomes in the 81,440 database).*

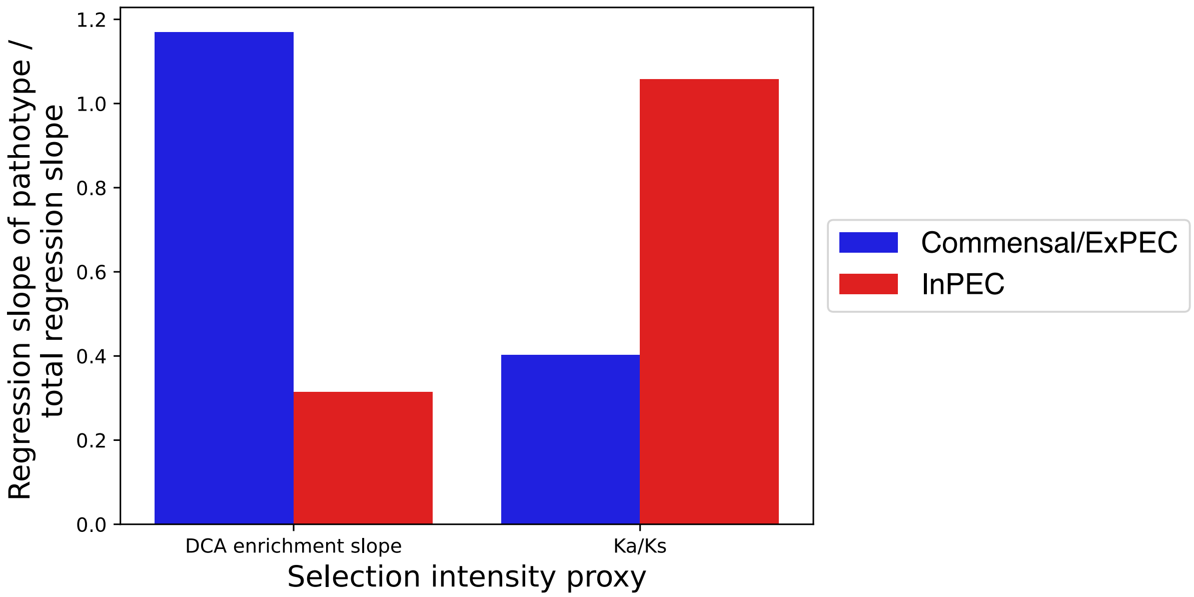


***Supplementary Fig. 9: Ratio of the regression slopes presented in Figure 9 by the regression slope computed across all clusters (whatever their pathotype), showing which parts of the clusters drive the regression for DCA enrichment slopes and for Ka/Ks.***
